# SPARROW reveals cell states and functions influenced by microenvironment zones in complex tissues

**DOI:** 10.1101/2024.04.05.588159

**Authors:** Peiyao A Zhao, Jessica Garber, Claire Gustafson, June Kim, Jocelin Malone, Adam Savage, Peter Skene, Xiao-jun Li

## Abstract

Spatially resolved transcriptomics technologies have significantly enhanced our ability to understand cellular characteristics within tissue contexts. However, they present a trade-off between spatial resolution and transcriptome coverage. This limitation, compounded with analytical tools treating cell type inference and cellular neighbourhood identification as separate processes, hinders a unified understanding of tissue features across scales. Our computational framework, SPARROW, infers cell types and delineates cellular organization patterns as microenvironment zones using an interconnected architecture. SPARROW algorithmically achieves single cell spatial resolution and whole transcriptome coverage by integrating spatially resolved transcriptomics and scRNA-seq data. Using SPARROW, we identified established and novel microenvironment zone-specific ligand-receptor mediated interactions in human tonsils, discoveries that would not be possible using either modality alone. Moreover, SPARROW uncovered novel cell states in the mouse hypothalamus, underscoring the influence of microenvironment zones on cell identities. Lastly, through its common latent spaces that facilitate cross-tissue comparisons, SPARROW revealed distinct inflammation states between different lymph node tissues. Overall, SPARROW integrates cellular gene expression with spatial organization, providing a comprehensive characterization of tissue features across scales and samples.

## Introduction

Spatially resolved transcriptomics (SRT) technologies have shown great promise in advancing our understanding of the organizational principles of diverse cell types within tissues during development and in disease progression [1, 2, 3, 4, 5, 6]. SRT techniques, however, present a trade-off between spatial resolution and transcriptomic coverage. High-resolution techniques like merFISH and seqFISH+ provide single-cell resolution but cover only a fraction of the transcriptome. Conversely, approaches like Visium perform whole-transcriptome measurements but at a reduced spatial resolution. Emerging methods such as Visium-HD and Slide-tag promise near single-cell resolution with comprehensive transcriptomic coverage, although they remain costly and are yet to be widely adopted [7]. This underscores the need for computational strategies that effectively bridge SRT with orthogonal whole transcriptome datasets, such as scRNA-seq data, to leverage spatial information gleaned from SRT for prediction of spatial localizations of cells in scRNA-seq data. Furthermore, many computational tools have been developed with the goal of performing cell type inference or spatial domain characterization, which are often treated as isolated tasks. Tools like Seurat, TANGRAM, and cell2location integrate SRT data with orthogonal scRNA-seq data to infer cell types without directly incorporating spatial context [8, 9, 10]. In contrast, a second class of methods such as BayeSpace, Giotto, SpaGCN, STAGATE, SpiceMIX and BANKSY integrate gene expression with spatial constraints to enable cell type identification or spatial domain delineation [11, 12, 13, 14, 15, 16]. The first category of algorithms, by disregarding spatial contexts, precludes the identification of spatially distinct cell states. The second category through convolution steps that merge spatially proximal transcriptomes tend to produce over-smoothed spatial domains that do not accurately reflect the complexity of underlying biology: even within the same spatial domain, distinct and biologically relevant cell neighbourhoods can exist. Moreover, the continuous nature of these cell neighbourhoods, characterized by gradual transitions from one to another, is not reflected in the designs of existing algorithms, limiting their ability to delineate finer changes to cell states in the context of the continuum of cell-to-cell interactions in the tissue space.

To this end, we developed SPARROW, a computational framework that performs integrative cell type inference and microenvironment zone delineation and is capable of predicting microenvironment zones for cells lacking spatial context such as those in scRNA-seq. We applied SPARROW to human immune tissues— including tonsils and lymph nodes (LNs)—and the mouse hypothalamus, all characterized by complex cell neighbourhoods that cannot be summarized by smooth spatial domains. In these diverse biological contexts, SPARROW uncovered novel cell states within the same cell types, attributed to their unique microenvironment zones. Furthermore, SPARROW identified known and novel microenvironment zone specific ligand-receptor (L-R) mediated interactions between cell types through both scRNA-seq based microenvironment zone predictions as well as direct geospatial modelling on SRT data. Lastly, SPARROW enabled comparative analysis of two LNs of distinct anatomical origins (mediastinal and mesenteric), revealing changes in cell type composition resulting in variations in tissue microenvironment zones that indicate differential inflammation states. Overall, we present SPARROW as a novel approach that fully integrates cellular gene expression with their spatial context, providing a comprehensive characterization of tissue features across scales and samples.

## Results

The first component of SPARROW, SPARROW-VAE (<u>v</u>ariational <u>a</u>uto<u>e</u>ncoder), constructs a shared latent space to co-embed the SRT expression matrix *X*_*N,M*_ and the orthogonal expression matrix such as scRNA-seq 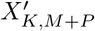 through a common latent representation *Z*, which encodes cell type information *L*. SPARROW-VAE also has the flexibility to perform cell type inference in the absence of 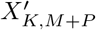, in which *Z* can be manually clustered and annotated. Subsequently, *Z* is ported into SPARROW-GAT (<u>g</u>raph <u>at</u>tention network), where *Z* serves as node features and spatial connectivity between neighboring cells serve as edges. SPARROW-GAT classifies microenvironment zones *H* that represents unique cellular neighborhoods formed by multiple cell types in a continuous space (**Figure 1A**). The trained SPARROW model has in turn learnt spatial relationships between cells in the SRT data *X* and can be further applied to 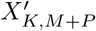 to perform microenvironment zone predictions for these cells that lack spatial localisation. This predictive capability extends spatial insights to cells that otherwise lack spatial context, thereby removing constraints imposed by gene panel designs on downstream analyses predicated on spatial colocalisation, such as the transcriptome-wide prediction of L-R interactions. Additionally, the construction of common latent spaces by SPARROW enables the projection of *X* and *H* from disparate experiments and conditions into a unified framework, allowing for direct comparisons.

**Figure 1:**
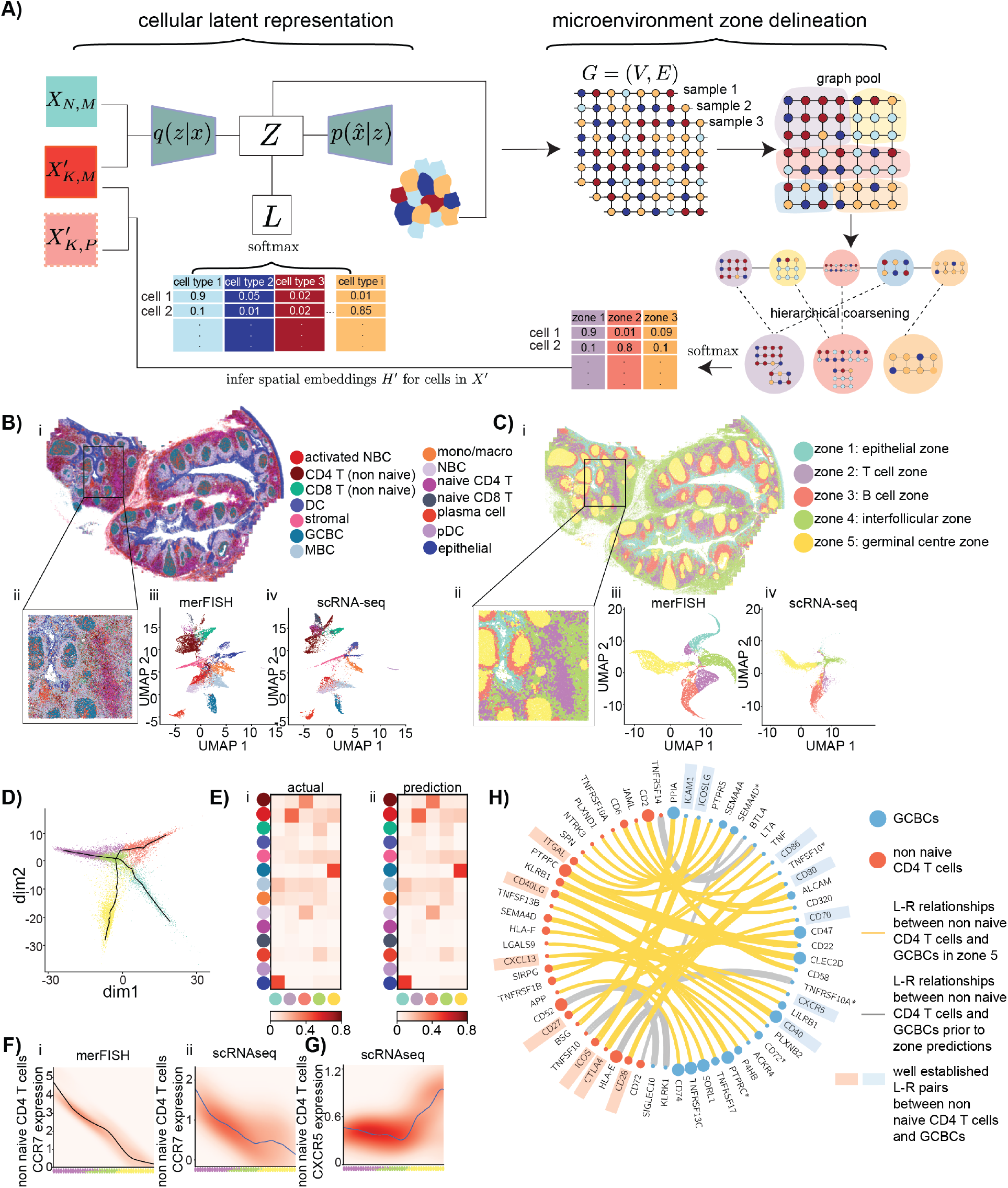
Integration of transcriptome and spatial context by SPARROW. **A**: Overview of SPARROW: the first component of SPARROW framework is a VAE for which the goal is to learn the latent representations of cells in SRT data that best preserve cell type specific information. Integer based SRT gene count matrices of shape *N* × *M*, where *N* denotes the number of cells and *M* denotes the number of genes in the gene panel covering partial transcriptome, and cell type labelled scRNA-seq count matrices of shape *K* × *M* are encoded as latent representation *Z*, the inference of which is achieved by minimizing KL divergence between the posterior *q*(*Z*| *X*), modelled as a multivariate Gaussian distribution, and the prior *p*(*Z*), while maximizing the log-likelihood 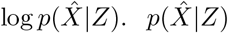 is modelled as a ZINB distribution, parameterized by *π*, the probability of excess zeros, *r*, the dispersion parameter and *ρ*, the rate of success. The parameterization of these distributions is implemented through neural networks (NNs). When cell type-labelled scRNA-seq data from the same tissue are available, *X*_*N,M*_ and 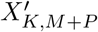, in which *P* denotes the number of genes not included in the gene panel, are co-embedded in the shared latent space, with scRNA-seq labels *L* modeled using categorical distributions parameterized by logit layers. To categorize the types of microenvironment zones present in the tissues, *Z* is integrated into a graph and ported into the SPARROW-GAT module. Here, *Z* serves as node features and an adjacency matrix *A*, which denotes the user defined k-nearest neighbors of cells in the tissue space, serves as edges. Through GAT operations, *Z* is output as a new latent embedding *H*, which is subsequently aggregated into distinct microenvironment zones through the minCUT algorithm. Furthermore, for cells in scRNA-seq data, whose *Z* is known but *A* is unknown, *A* can be deduced from SRT data, leveraging latent space *Z* similarities between scRNA-seq and SRT cells as *X*_*N,M*_ and 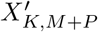 are co-embedded. This inference enables SPARROW-GAT to perform microenvironment zone prediction for scRNA-seq cells. See Methods for detailed formulation. **B**: Cell type inference by SPARROW-VAE in tonsil EXP429 by co-embedding merFISH data *X* and scRNA-seq data *X*^′^: inferred cell types in the tonsil section EXP429 (i). A 2000 pixel × 2000 pixel window of the tissue section was zoomed in for details of cell type localization (ii). 2D UMAP visualization of *Z* showing cells in tonsil merFISH data *X* (iii) and scRNA-seq data *X*^′^ (iv) sharing the same latent space and forming distinct clusters informed by cell types. **C**: Microenvironment zone assignments and predictions in merFISH data of tonsil EXP429 and tonsil scRNA-seq data, respectively: microenvironment zone assignments in the tonsil section EXP429 (i). A 2000 pixel × 2000 pixel window of the tissue section was zoomed in for details of the distinct spatial localization patterns of zones (ii). 2D UMAP visualization shows microenvironment zone assignment *H* of cells in tonsil merFISH data (iii) and microenvironment zone predictions *H*^′^ in tonsil scRNA-seq data (iv) in the shared latent space and forming distinct clusters informed by zones. The colour-coding is consistent between **C, D** and **E. D**: A principal graph was computed for *H* of merFISH data of tonsil EXP429. Cells projected onto the graph space are represented by dots colored according to their respective microenvironment zone assignment. Cells in zones 1-5 occupied continuous yet non-overlapping niches in the graph space. Branches in the principal graph represent the spatial relationships between microenvironment zones within the tissue. **E**: Heatmaps showing cell type composition of microenvironment zones in merFISH data of tonsil EXP409 (**i**) and predicted microenvironment zones based on tonsil scRNA-seq (**ii**). The rows indicate cell types and add up to 1 and the columns indicate microenvironment zones. **F**: CCR7 expression of non naive CD4 T cells along the selected trajectory as shown in dark blue in Supplementary Figure 9C, which traverses zone 2, the T cell zone, through zone 4, the interfollicular zone and ending in zone 5, the GC zone, of the principle graph computed using either *H* of tonsil merFISH (**i**) or *H*^′^ prediction from tonsil scRNA-seq data (**ii**). Below the x-axis, pie charts display the percentage of cells with colour-coded zones projected along the selected trajectory in the graph. **G**: CXCR5 expression of non naive CD4 T cells along the selected trajectory as in **F** of the principle graph computed using *H*^′^ prediction from tonsil scRNA-seq data. **H**: A Circos plot showing significant L-R pairs between non naive CD4 T cells and GCBCs in zone 5, the GC zones (yellow) or regardless of zone assignments (gray), identified using cellphonedb modelling. Significance was defined as p values *<* 0.05. The widths of connectors are positively correlated with interaction scores. Nodes represent genes expressed in either non naive CD4 T cells (red) or GCBCs (blue), with node sizes reflecting the expression percentage in their associated cell types. Known L-R pairs are highlighted in red-blue block pairs. For genes expressed in both cell types, those in GCBCs are distinguished with asterisks.

To demonstrate the capability of SPARROW in integrating SRT *X*_*N,M*_ and orthogonal scRNA-seq 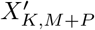, we applied SPARROW to merFISH data of a human tonsil section (EXP429). *X*_*N,M*_ from EXP429 was fed into SPARROW-VAE and co-embedded with *X*^′^ from a published and annotated tonsil scRNA-seq dataset [17] (**Supplementary Figure 1A**,**B**) resulting in 14 cell types forming distinct clusters in the shared common latent space (**Figure 1B, Supplementary Figures 1C**,**D, 2**). The accuracy of SPARROW-VAE outperformed state-of-the-art cell inference methods TANGRAM and SPICEMIX (**Supplementary Figure 3**), as validated using both simulated data (**Supplementary Figure 4**) and orthogonal multiplexed immunofluorescence protein imaging (mxIF) (**Supplementary Figure 5**). We next applied SPARROW-GAT to *Z* of EXP429 tonsil to algorithmically characterize tissue microenvironment zones (**Supplementary Figure 6A**,**B**) and identified 5 zones occupying distinct spatial niches both in tissue space and latent space (**Figure 1C**). Moreover, they are characterized by unique spatial interaction patterns and distinctive cell type composition (**Supplementary Figure 6C**,**D**), aligning with biological expectation. Briefly, zone 1, the epithelial zone, was primarily composed of interacting epithelial cells and memory B cells (MBCs) (**Supplementary Figure 6Di**). Zones 2 and 4, representing different types of zones within the tonsil paracortex, were identified as the T cell zone and the interfollicular zone, respectively (**Supplementary Figure 6Dii**,**iv**). Zone 3, the B cell zone, encircled germinal centres (GCs) and mainly consisted of interacting naive B cells (NBCs), activated NBCs and MBCs (**Supplementary Figure 6Diii**). Lastly, zone 5, the GC zone, was primarily composed of GC B cells (GCBCs) (**Supplementary Figure 6Dv**). In addition to capturing static cell interaction patterns, *H* captures continuous transitions between microenvironment zones based on cellular neighbourhood changes (**Figure 1D**). For instance, zone 2, the T cell zone, is connected to zone 3, the B cell zone via zone 4, the interfollicular zone, which is composed of a mixture of T and B cells (**Figure 1D, Supplementary Figure 6D**). These trajectories recapitulate known dynamics of T-B cell migration out of their respective zones and subsequent interactions in the interfollicular zone. We next applied the SPARROW-VAE and -GAT models trained on EXP429 to three additional tonsil sections and observed consistent cell type composition and localisation patterns of microenvironment zones across sections (**Supplementary Figures 7**,**8**) highlighting the efficacy of latent space to unify samples.

The integrative framework of SPARROW further enables the prediction of microenvironment zone latent embedding *H*^′^ for cells in scRNA-seq data 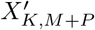, thus computationally achieving single cell spatial resolution with whole transcriptome data. To assess prediction accuracy, we first benchmarked *H*^′^, predicted using only the expression matrix of cells in a test tonsil tissue (EXP501), against the actual microenvironment zone assignments *H* inferred using both the expression matrix and spatial adjacency, and showed that prediction accuracies reaching approximately 80% (**Supplementary Figure 9A**). Furthermore, cell type composition of predicted zones showed high concordance with actual zone assignments (**Figure 1E**). We further predicted *H*^′^ for cells in tonsil scRNA-seq. The quality of this prediction is supported by the highly similar gene specificity patterns across both the known zones of tonsil merFISH data *X*_*N,M*_ and the predicted zones in scRNA-seq 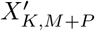 (**Supplementary Figure 9B**). Cells’ positions along trajectories presented in **Figure 1D** hold important information regarding cell state transitions across various microenvironment zones, even within the same cell type. Using SPARROW’s zone predictions for cells in 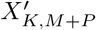, we can identify expression changes of genes absent in the SRT gene panel in cells along the trajectory, thus revealing cell identities that couldn’t be otherwise confirmed using *X* alone. Specifically, focusing on a selected path starting in zone 2, the T cell zone, traversing through zone 4, the interfollicular zone, and ending in zone 5, the GC zone, non naive CD4 T cells exhibit a bimodal distribution with predominant presence along zone 2 and zone 4, with a small population along zone 5, hypothesized to be follicular helper T cells, a label absent from the original *X*^′^ (**Supplementary Figure 9C**). Further querying of transcriptomic variations of non naive CD4 T cells along this path revealed a progressive reduction in CCR7 expression—a gene downregulated as CD4 T cells migrate from the T cell zone to the GC zone [18]—both in observed merFISH data (**Figure 1Fi**) and scRNA-seq based zone predictions (**Figure 1Fii**). Concurrently, genes absent from the merFISH gene panel, including CXCR5 and IL21, which are markers for follicular helper CD4 T cells, along with other novel markers, were predicted to increase in expression as CD4 T cells migrate into zone 5 (**Figure 1G, Supplementary Figure 9D**,**E**). This has confirmed the identity of the subpopulation of non naive CD4 T cells residing in GCs as follicular helper CD4 T cells, highlighting SPARROW’s predictive ability in surpassing the limitations of partial gene panels and leveraging the complete transcriptome to reveal dynamic states of cells within tissue spaces.

The predictive framework of SPARROW further facilitates the delineation of transcriptome wide L-R interactions between specific cell types in a microenvironment zone aware fashion. We performed L-R analyses using CellPhoneDB [19], either confining the analyses to cells predicted to occupy the same microenvironment zone or without consideration for zone predictions. Focusing on CD4 T cells and GCBCs predicted to be in zone 5, the GC zones, the former zone aware approach identified established L-R relationships including CD40LG-CD40, ICOSLG-ICOS and CTLA4-CD80, which are known to be important in T-B cell cross talk in GCs [20] (**Figure 1H**). In contrast, the latter zone unaware approach failed to identify these established relationships (**Figure 1H**). The same pattern holds true for L-R relationships between other cell types and zones (**Supplementary Figure 10**). Furthermore, this zone aware approach also uncovered novel L-R pairs, thereby opening up new grounds for future hypothesis testing.

To demonstrate SPARROW’s ability to delineate complex and unresolved cell neighbourhoods and identify novel cell states influenced by microenvironment zones, we applied SPARROW to merFISH data from serial sections of a mouse hypothalamus ([21]), a region characterised by complex neighbourhoods that cannot be simplified into homogeneous spatial domains. SPARROW projected sections into a unified latent space, revealing 11 microenvironment zones, each characterized by distinct cell compositions and neighbourhood patterns (**Figure 2A**,**B, Supplementary Figures 11, 12**). To demonstrate how microenvironment zones can impact cell states within the same cell type, we focused on a specialised cell type, SCH Six6 Cdc14a Gaba, which occurs exclusively in the suprachiasmatic nucleus (SCH) and are implicated in the regulation of the circadian clock ([22]). These cells are primarily assigned to zones 2 and 6 (**Figure 2C**). A comparative analysis revealed that SCH Six6 Cdc14a Gaba cells in zone 6 contact oligodendrocyte progenitor cells (OPCs) with significantly higher frequency than those in zone 2 (**Figure 2D**). SCH Six6 Cdc14a Gaba cells in zone 2 and zone 6 are also molecularly different, supported by differential gene expression patterns between the two groups (**Figure 2Ei,ii**). Upregulated genes in SCH Six6 Cdc14a Gaba in zone 6 were specifically enriched for GABAergic and Glutamatergic synapse pathways (**Figure 2Eiii**), which, together with increased interaction with OPCs, aligns with previous findings suggesting neuronal activities’ modulation of OPCs ([22]). Overall, this supports SPARROW’s capability in categorising intricate local neighbourhoods and in differentiating fine cell states that can be explained by their microenvironment zones, allowing for the generation of new hypotheses regarding cellular identities within spatial contexts.

**Figure 2:**
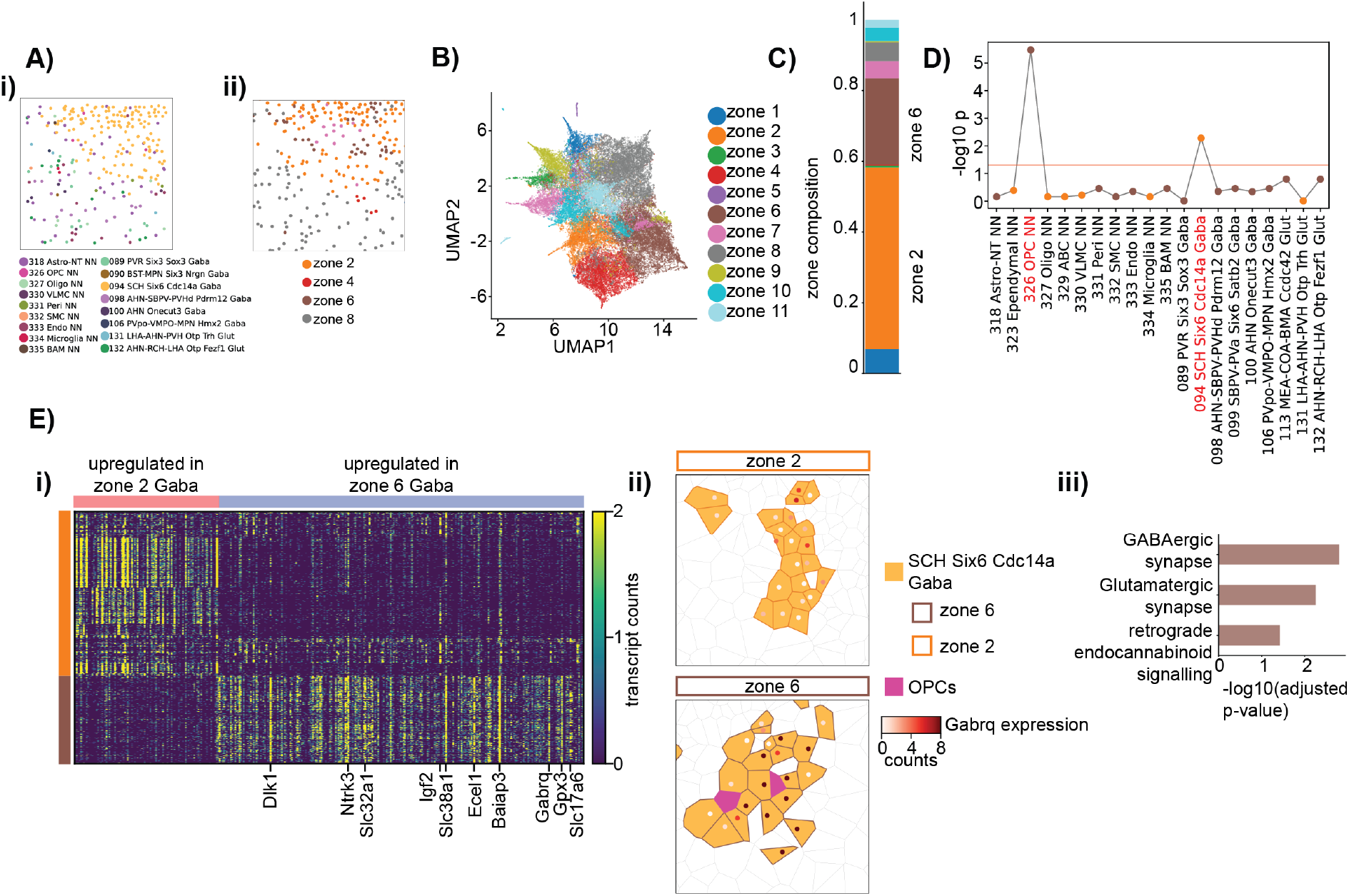
SPARROW reveals microenvironment zone specific cell states in the mouse hypothalamus. **A**: Spatial visualisation of cells in a section of the mouse hypothalamus slide [21], colour-coded by cell type subclass according to the colour scheme of the original publication (**i**) and their microenvironment zone assignments (**ii**). **B**: 2D UMAP visualization of microenvironment zone latent embedding *H* showing cluster separation. **C**: A bar plot showing zone assignments for SCH Six6 Cdc14a Gaba cells in the hypothalamus. 51% and 24% are assigned to zone 2 and zone 6, respectively. **D**: A plot showing Benjamini-Hochberg corrected -log10 p values for contact frequency differences between cell types shown on the x-axis and SCH Six6 Cdc14a Gaba cells in zones 6 versus zone 2 using a chi-squared test. The red line indicates significance threshold of (-log10(0.05)). Cell types showing more frequent contact with SCH Six6 Cdc14a Gaba cells in zones 6 than in zone 2 are shown as brown dots while those that show opposite are shown as yellow dots. OPC interactions show the most significant difference and are increased in zone 6. **E**: **i**: A heatmap showing expression of genes that are significantly different between zone 2 and zone 6 Six6 Cdc14a Gaba cells. Cells and genes correspond to rows and columns respectively. Rows are sorted such that zone 2 Six6 Cdc14a Gaba cells are on top (indicated by the orange bar). Columns are sorted such that genes that are upregulated in zone 2 Six6 Cdc14a Gaba cells are shown on the left (indicated by the red bar). Top 10 genes that are most significantly upregulated in zone 6 Six6 Cdc14a Gaba cells are labelled on the x-axis. **ii**: Representative Voronoi diagrams showing SCH Six6 Cdc14a Gaba cells in zone 2 (outlined by orange lines) and zone 6 (outlined by brown lines). SCH Six6 Cdc14a Gaba cells in zone 6 are in contact with OPCs (shown as pink polygons). Gabrq, whose expression is upregulated in zone 6 SCH Six6 Cdc14a Gaba cells, is shown as colour coded dots. The colour gradient is positively correlated with transcript counts. Cells assigned to other zones are white polygons outlined with black lines. **iii**: Gene set enrichment analysis (GSEA) showing KEGG pathways that are significantly enriched in upregulated genes in zone 6 Six6 Cdc14a Gaba cells. GSEA analysis using upregulated genes in zone 2 Six6 Cdc14a Gaba cells did not return significant KEGG pathways.

Lastly, SPARROW enables comparisons across tissue sections of similar but not identical origins through its common latent spaces, revealing important alterations in tissue states. We trained SPARROW-VAE and -GAT on merFISH data generated on a human mediastinal LN (EXP621) and further applied trained models to a mesenteric LN section (EXP505) of the same donor origin (**Figure 3A**,**Supplementary Figures 13, 14, 15**). We observed expansion and reduction in specific cell populations with corresponding microenvironment zone changes (**Figure 3B, Supplementary Figures 16A**,**B**). Notably, zone 2 and 5, showed marked enrichment, in the mediastinal LN (EXP621) compared to the mesenteric LN (EXP505), consistent with increases in their main components, GCBCs and CD4 T cells, respectively (**Figure 3B**). This is indicative of increased inflammatory activity in EXP621 compared to the EXP505 [23]. To further query this, we examined if EXP621 showed microenvironment zone specific L-R interactions characteristic of inflammation. We focused on a well characterized L-R pair CD40LG-CD40 whose interaction plays an important role in T-B cell cross-talk in lymphoid follicles formation and is elevated in inflammation [24]. In EXP621, we observed statistically significant colocalisation at distances of 2-3 median cell radii, the range consistent with cell-to-cell interaction in zone 2, the follicular zone, and to a lesser extent, zone 3, the B cell zone, but not in the remaining zones or without considering zone assignments, highlighting the inflammation related L-R interaction being spatially localised in defined microenvironment zones (**Figure 3Ci, Supplementary Figures 17, 18**). This zone dependent CD40LG-CD40 interaction was found not to be significant in EXP505, consistent with reduced inflammation (**Figure 3Cii, Supplementary Figure 18**). Overall, this highlights SPARROW’s capacity to delineate and compare spatially distinct biological processes across samples, thereby yielding insights into variations in tissue states.

**Figure 3:**
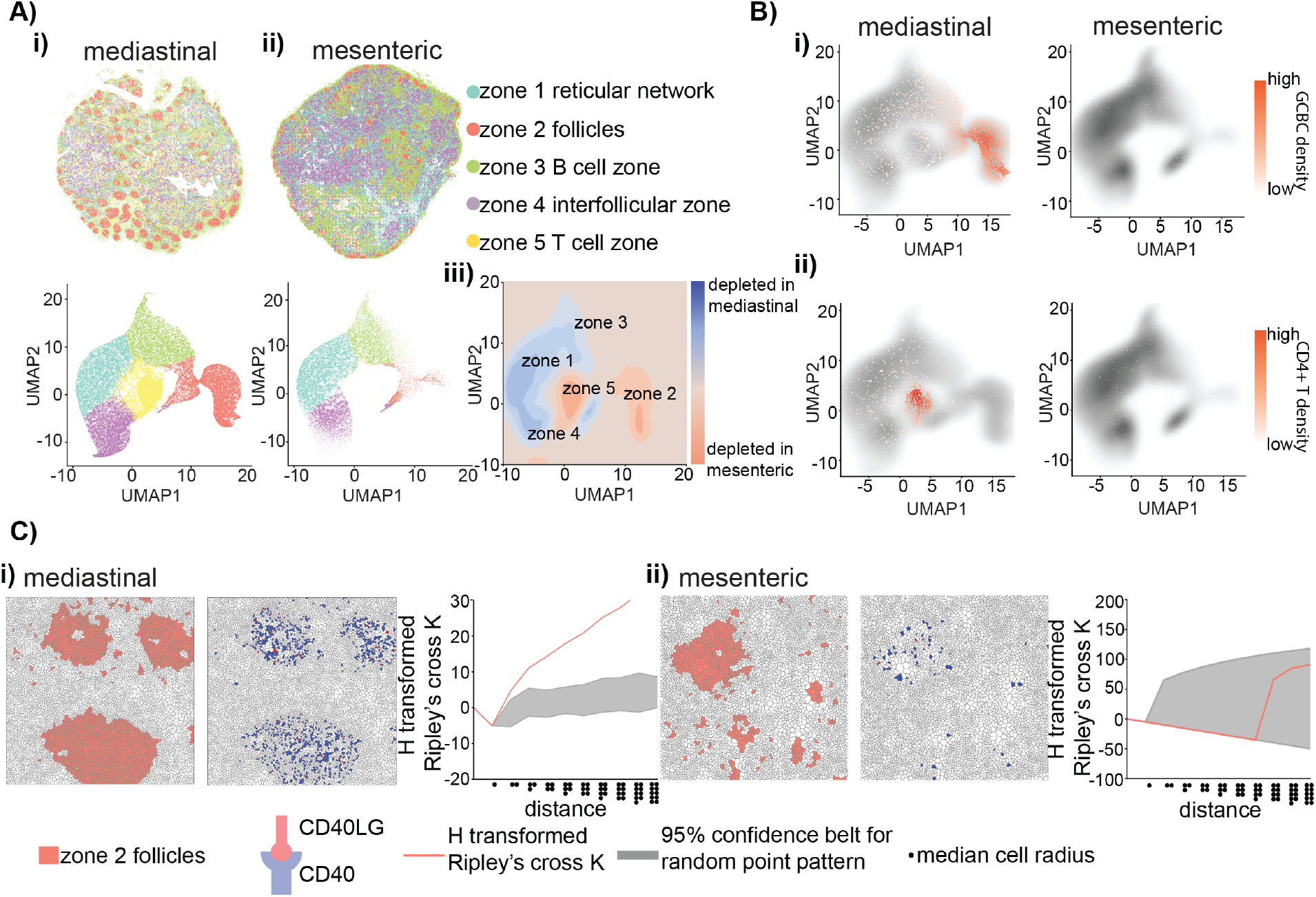
SPARROW unveils changes in tissue states between a mediastinal LN and a mesenteric LN. **A**: Spatial visualisation of cells in the mediastinal LN (EXP621) (**i**) and mesenteric (EXP505) LN (**ii**), colour-coded by their microenvironment zone assignments (top row). 2D UMAP visualisation of microenvironment zone latent embedding *H* is shown on the bottom row. **iii**: 2D UMAP visualization of differentials in microenvironment zone density between the mediastinal LN and the mesenteric LN. **B**: Density plots of *H* of the mediastinal (EXP621) and mesenteric (EXP505) LN showing microenvironment zone distribution differences. Density of GCBC cells and non naive CD4 T cells, the main constituents of zone 2 and 5 respectively, are shown in red. **C**: Ripley’s K calculation showing significant zone 2 specific colocalisation between CD40 positive (receptor, blue) and CD40LG positive (ligand, red) cells in the mediastinal LN (**i**) but not in the mesenteric LN (**ii**). Zone assignments (left) and co-localisation patterns (middle) of CD40 positive (receptor, blue) and CD40LG positive (ligand, red) cells are represented as Voronoi diagrams. H-transformed Ripley’s cross K of CD40 positive cells relative to CD40LG positive cells as a function of increasing distances (black line) in units of median cell radii is shown in the right-most panel. 1000 iterations of spatial random patterns were generated by randomly shuffling CD40 positive and CD40LG positive cell labels and Ripley’s cross K was calculated for every pattern iteration. The 95% confidence envelope of the random patterns was plotted (gray belt) and compared against the experimental curve. The experimental curve being above the confidence belt indicates significant colocalisation at the distances indicated on the x-axis.

## Discussion

SPARROW is a computational framework designed for analysing single cell resolution SRT data that integrates cellular gene expression and spatial organization. It outperformed two leading methods in cell typing accuracy and effectively delineated complex tissue microenvironment zones corresponding to units of biological functions, beyond smooth spatial domains typically defined by existing SRT methods. By integrating SRT and scRNA-seq in the latent space, SPARROW computationally achieves what is experimentally challenging: single-cell spatial resolution across whole transcriptomes. We showed that such computational inference facilitated the discovery of novel L-R mediated cell interactions within specific microenvironment zones. Furthermore, the integrative formulation has enabled SPARROW to identify novel cell states, informed by interactions with diverse cell types in different microenvironment zones. Lastly, SPARROW’s latent embedding spaces enabled direct comparisons across tissue samples and generated insights into the changing tissue states.

While in this manuscript SPARROW has been applied to merFISH data for biological discoveries, it is applicable to any other types of single cell resolution SRT data. Moreover, given the continuous evolution of SRT techniques and the addition of multimodal aspects to spatial data (reviewed in [25]), SPARROW can be readily adapted to encode additional data modalities and thereby shed new light on the interplay between RNA-protein or epigenome-RNA within tissue microenvironment zones.

## Methods

### LN and tonsil tissue collection

LN and tonsil tissues were collected post-mortem by the National Disease Research Interchange (NDRI). NDRI maintains a Federal Wide Assurance (FWA00006180) agreement with the DHHS, Office for Human Research Protections to comply with federal regulations concerning research involving human subjects. These studies were done in accordance with the Declaration of Helsinki and were approved by Allen Institute for Immunology Internal Review Board.

### merFISH data generation

5*μ*m sections were cut from human LN and tonsil Formalin-Fixed Paraffin-Embedded (FFPE) blocks. Sections were floated in an RNAseq free, 41°C water bath and adhered to proprietary MERSCOPE slides. Sectioned tonsil and LN samples were processed in accordance with the Vizgen Formalin-Fixed Paraffin-Embedded Tissue Sample preparation protocol. Cell boundaries were stained using Vizgen’s Primary and Secondary antibody mix (Vizgen Cell Boundary Primary Stain Mix PN # 20300010 and Cell Boundary Secondary Stain Mix PN # 20300011). For tissue clearing, the samples were treated with a digestion mix before clearing up to 24 hours in a humidified 47C cell culture incubator then transferring to a 37C cell culture incubator for up to 4 days. Once the tissue became transparent, autofluorescence quenching was undertaken using the MERSCOPE Photobleacher (PN # 10100003) and incubating at room temperature for at least three hours. merFISH probe hybridization was completed with 400 (tonsil) and 130 (LN) -gene custom panels (PN # 10400001 or PN # 10400003, respectively). Cell segmentation was performed on merFISH instrument using CellPose algorithm with cell-boundary stain 3 staining as input.

### merFISH data preprocessing

For tonsil and LN merFISH data, cells with transcript counts in the lower and highest 5% were removed from downstream analyses for they likely represent technical artifacts. Transcripts within the gene panel that show extremely low variance or have unusually low or high counts (thresholds to be determined by the user) may also be removed. The merFISH data are formatted into parquet files which contain spatial coordinates indicating cell boundaries and centroids and transcript counts for cells.

### SPARROW formulation

SPARROW is composed of two connected formulations: a VAE for cell type inference and a GAT for microenvironment zone delineation.

#### VAE

The VAE is composed of an encoder and a decoder. The original formulation was proposed in [26] and has been adapted for single cell data analysis by several tools published to data, including scVI, scVAE and bmVAE [27, 28, 29], among others.

#### Encoder

Let 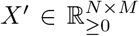 be the scRNA-seq cell by gene count matrix of shape *N* × *M* for which *N* denotes the number of cells and *M* the number of genes. Let 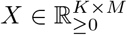 be the ST cell by gene count matrix of shape *K* ×*M*, for which *K* denotes the number of cells and *M* is the number of genes. The encoder encodes *X* and *X*^′^ over the latent space *Z* using fully connected NNs with a user defined number of hidden layers to parameterize a posterior probability distribution *q*(*Z*| *X*). *q*(*Z*| *X*) is assumed to be a multivariate gaussian distribution with a mean vector *μ*(*X*) and a diagonal covariance matrix Σ(*X*). The latent representation *Z* is sampled from the multivariate Gaussian distribution *Z* ∼𝒩(*μ*(*X*), Σ(*X*)). In the case that labelled scRNA-seq data with cell labels *L* is used, *L* follows a categorical distribution parameterized by the logits, which is sampled from the posterior distribution *q*(*logits*|*Z*).

#### Decoder

The latent representation *Z* is transformed by the decoder to reconstruct input data *X* as 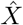 in a series of operations that mirror that of the encoder and aim to parameterize the distribution *p*(*X* |*Z*). *p*(*X*| *Z*) is assumed to follow a zero-inflated negative binomial (ZINB) distribution due to the data characteristics of ST data. The probability mass function (PMF) for ZINB is as follows:

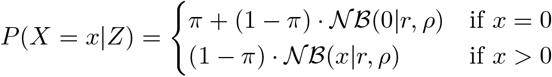

where *π* is the probability of an excess zero and 𝒩ℬ(*x r, ρ*) is the PMF of the Negative Binomial distribution where r is the dispersion parameter and *ρ* is the probability of success. *π* and *ρ* are in turn derived from a multivariate Gaussian distribution parameterized by NNs in the decoder and *r* is set as a hyperparameter.

#### loss function

When running in the unsupervised mode, the loss function for the VAE module is evidence lower bound (ELBO), which is composed of two components: the reconstruction loss and the regularization term. The reconstruction loss represents the difference between the observed data *X* and the decoded data 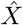 and is defined as:

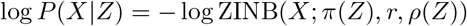

The regularization term accounts for the KL divergence between the encoded distribution and the prior distribution *Q*(*Z* |*X*) and the prior distribution *P* (*Z*). This term minimizes the difference between the encoded distribution and the prior distribution:

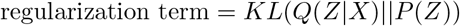

In the supervised mode where labelled scRNA-seq *X* is co-embedded with *X*, a third loss term, the classification loss, is added:

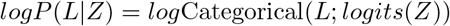

where logits(Z) are the logits derived from Z for the categorical distribution from which *L* is sampled. In summary, the loss function is as follows:

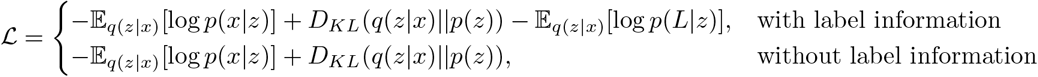

Loss minimization was implemented using stochastic variational inference (SVI) in the Pyro framework (https://github.com/pyro-ppl/pyro). Shapley additive explanations (SHAP) scores [30] were used to find top contributing feature genes for SPARROW-VAE cell type classification.

#### GAT

The latent representation *Z* from SPARROW-VAE is integrated into an undirected graph 𝒢(𝒱, *ℰ*) in which 𝒱 is the collection of cells as nodes with *Z* being node features and *ℰ* is the spatial neighborhood of cells expressed as an adjacency matrix *A* in which *A*_*i,j*_ = 1 if cells *i* and *j* are the k nearest neighbors to each other. We defined k=7 for our tonsil and LN datasets and k=5 for the mouse hypothalamus dataset from [21] due to its smaller cell population in comparison to human datasets. SPARROW-GAT module uses the pytorch implementation of the GAT formulation proposed in [31] and computes the new node representation 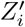 for node *i* in the *h*-th head as follows:

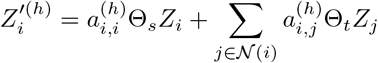

where 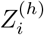 is the input node feature for node *i*, 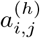 is the learnable attention coefficient between node *i* and its neighbor node *j* for the *h*-th head, Θ_*s*_ and Θ_*t*_ are learnable weight matrices for source and target nodes, respectively, and 𝒩(*i*) is the neighborhood of node *i*. For tonsil and LN datasets, the number of attention heads was chosen to be 3. For the mouse hypothalamus dataset, the number of attention heads was increased to 6 due to more cell types present. 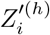 is followed by an exponential linear unit (ELU) activation and dropout regularization. Finally, a linear transformation is applied to output node embeddings *H* of dimension *N* × *C* where *N* represents the number of nodes and *C* denotes the number of user-defined pooled graphs, i.e. microenvironment zones.

#### loss function

*H* is partitioned using MinCUT algorithm as defined in [32]. Briefly, MinCut aims to find optimal partitions by minimizing the sum of two loss terms: MinCut loss and orthogonality loss. MinCut loss can be calculated as such:

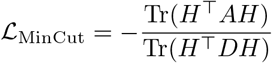

where *H* is the node latent embedding output from GAT, *A* is the adjacency matrix and *D* is the diagonal degree matrix. *Tr* indicates the trace operation. This loss term is designed to minimize the connections between partitions and maximize those within partitions. Orthogonality loss is defined as:

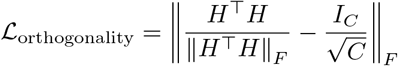

where *I*_*C*_ is the identity matrix of size *C*, the number of clusters and ∥·∥ _*F*_ is the Frobenius norm. This loss term penalizes degenerate solutions where all nodes are assigned to the singular partition. The loss function was optimized using ADAM [33], a first-order gradient based stochastic optimizer, implemented in pytorch. Training convergence is user defined and was set at ℒ *<* 0.01 for the trained models used in this manuscript.

### *H*^′^ prediction for cells lacking edge information

For cells in scRNA-seq data, *Z* is known whereas *A* needs to be inferred by leveraging latent space *Z* similarities between scRNA-seq and SRT cells as *X*_*N,M*_ and 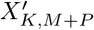 are co-embedded by SPARROW-VAE. Specifically, for cell *i* in scRNA-seq, top n closest points in the SRT data are selected based on the Euclidean distance in *Z* such that 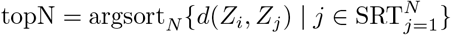. Subsequently, the adjacency relationships for each identified cell *j* in the SRT data, represented by *A*_*j,K*_, signifying cell *j* and its K nearest neighbours in the SRT tissue space. For each cell *k* in the K nearest neighbours, we in turn seek the nearest cells in the scRNA-seq in latent space *Z*, thereby inferring edges from the original cell *i* in scRNA-seq, 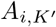, such that 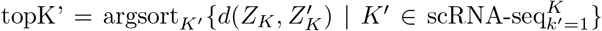. Edges 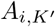 can then be ported along with node features *Z* into SPARROW-GAT to infer microenvironment zones for cells in scRNA-seq data.

### Simulation for benchmarking

N cells were randomly selected for each cell type in three separate fields of view in EXP409. For cell types identified by both SPARROW and SPICEMIX, only cells with labels agreed upon by both methods were selected. For cell types not identified by SPICEMIX, SPARROW labels were used. Cells were associated with spatial coordinates and expression vectors. Expression was used for SPARsim [34] parameter estimation using the command `SPARSim_estimate_parameter_from_data`. Estimated parameters were used for generating simulated data with the command `SPARSim_simulation`. Each simulated cell type was assigned spatial coordinates to mimic their spatial distribution in the real data. Specifically, a kernel density estimation (KDE) was fitted to the spatial coordinates of cells of a given cell type to model their spatial distribution. New coordinates were drawn randomly from this density function.

### mxIF imaging using Cell DIVE

#### Slide preparation and blocking

A 5*μ*M FFPE tonsil tissue slide close to that which was used for EXP409 was de-paraffinized using xylene and ethanol gradations, then rehydrated in 1x PBS. Tissue was permeabilized for 10 minutes with 0.3% Triton X-100 (Promega, H5141) and washed in 1x PBS. Heat Induced Epitope Retrieval (HIER) was performed with NxGen Decloaking Chamber™(Biocare, DC 2012) in successive baths of 110° Citrate pH 6 (Vector Laboratories, H-3300-250) and 85-110° Tris-EDTA pH 9 (VWR, TS22803-1000 and Sigma, E5131-100G) for 20 minutes each. Tissue was blocked in 10% reconstituted donkey serum (Jackson, 017-000-121), 3% bovine serum albumin (Sigma, A2153-50g) in 1x PBS. Tissue was washed in 1x PBS, then stained with 100 *μ*g/mL DAPI (Thermo Fisher, D3571) for 15 minutes. Slides were loaded into ClickWell® plates (Leica Microsystems, 29626840) for coverslip-free staining and imaging. For imaging, ClickWells® were filled with 0.8% propyl gallate (Sigma, P3130-100G), 50% glycerol (Sigma #G5516-500ML) in 1x PBS mounting media. For storage, ClickWells® were filled with 50% glycerol in 1x PBS storage media.

#### ROI selection and autofluorescence imaging

The Leica Cell DIVE was used for all immunofluorescence imaging. Whole tissue images were collected at 2x, 10x, and 20x. The 10x virtual H&E image was used to identify and select regions of interest. Autofluorescence imaging of whole tissue and selected ROIs were collected and retained for spectral subtraction from biomarker imaging.

#### Biomarker staining, imaging, and dye inactivation

Multiplex imaging was performed in 3 rounds of immunostaining and dye inactivation. Tissue was stained with CD4, CD68, Ki-67, CD8a, CD20 and CD3d. Each round of staining was a mix of 2-4 antibodies in 3% BSA in 1X PBS. Each antibody was directly conjugated to a different channel of fluorophore (AF488, AF555, AF647, AF750) to allow for simultaneous staining and spectrally distinct signal during imaging. Following autofluorescence imaging, mounting media was removed from the tissue/ClickWell® system using washes of 0.01% Tween 20 (Sigma, P1379-100mL) in 1X PBS. Antibody and 3% BSA was manually pipetted into the tissue/ClickWell® system and incubated on the benchtop at room temperature for 1 hour. After removing the antibody mixture with washes of 0.01% Tween, the slide was stained with 100 μg/mL DAPI, washed with 0.01% Tween, and the ClickWell chamber was filled with 0.8% propyl gallate/50% glycerol. 20X biomarker imaging was performed, and the Cell DIVE system used previously collected autofluorescence images to spectrally subtract background fluorescence from 20x biomarker images. Following imaging, mounting media was washed with 0.01% Tween, and fluorophores were bleached for 15 minutes using 0.1M NaHCO_3_ pH 11 (Sigma, S5761-500G) and 3% H_2_O_2_ (Sigma, H1009-500ML). Tissue was stained with 100 *μ*g/mL DAPI for 2 minutes and washed with 0.01% Tween, then returned to 0.8% propyl gallate/50% glycerol and autofluorescence images were collected for the next round of spectral subtraction. This was repeated a total of 3 times for a total of 9 biomarkers.

#### Cell DIVE image processing

Multiplexed images were stitched, registered, and aligned by the commercial Cell DIVE software. Image layering and biomarker thresholding was performed in HALO® Imaging Software (Indica Labs). Segmentation was performed using the built-in deep learning cell segmentation module within HALO software (https://indicalab.com/halo/) with domain expert guided manual boundary correction. Phenotyping for segmented cells was performed manually by domain experts based on the following logic:

| CD4 | CD68 | Ki-67 | CD8a | CD20 | CD3d | Cell Type |
| --- | --- | --- | --- | --- | --- | --- |
| + |  |  |  |  | + | CD4 T |
|  |  |  | + |  | + | CD8 T |
|  |  |  |  | + |  | non-GCBC B |
|  |  | + |  | + |  | GCBC |
|  | + |  |  |  |  | monocytes |

### Spatial trajectory analyses

Principal graphs were computed for node embeddings *H* using the algorithm proposed in [35] and its python implementation [36, 37]. Number of principal points *P* was decided using the formula presented in [36]. A cell in *H, h* ∈ *H* was projected to the nearest principal point *p* ∈ *P* according to 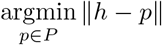 in which ∥*h* − *p*∥ represents the euclidean distance between *h* and *p*.

### Ripley’s cross K calculation

Let *R* = {*r*_1_, *r*_2_, …, *r*_*N*_} be all receptor positive cells and *L* = {*l*_1_, *l*_2_, …, *l*_*K*_} be all ligand positive cells. Ripley’s cross K measures if the number of *l ∈ L* within a certain distance from of *R* is higher than expected if *R* and *L* were independently and randomly distributed. Formally, Ripley’s cross K for *R* and *L* at the distance *d* is defined as follows:

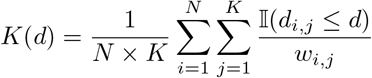

*w*_*i,j*_ is an boundary effect correction factor to account for the fact that cell *l* close to the edge of the defined area has a lower change of finding a neighboring cell *r* within *d. w*_*i,j*_ was calculated as follows:

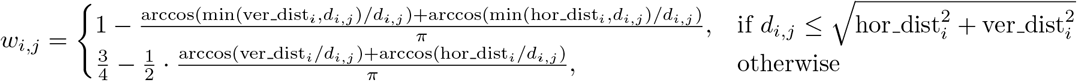

where hor_dist_*i*_ and ver_dist_*i*_ are the minimum horizontal and vertical distances from cell *i* to the edge of the user-defined area of interest.

The confidence belt was generated with the Monte Carlo method by shuffling the labels between *R* and *L* for 1000 iterations.

## Code Availability

The source code of SPARROW and test data can be accessed at https://github.com/peiyaozhao617/SPARROW.

