## Supplementary Figures for "SPARROW reveals cell states and functions influenced by microenvironment zones in complex tissues"

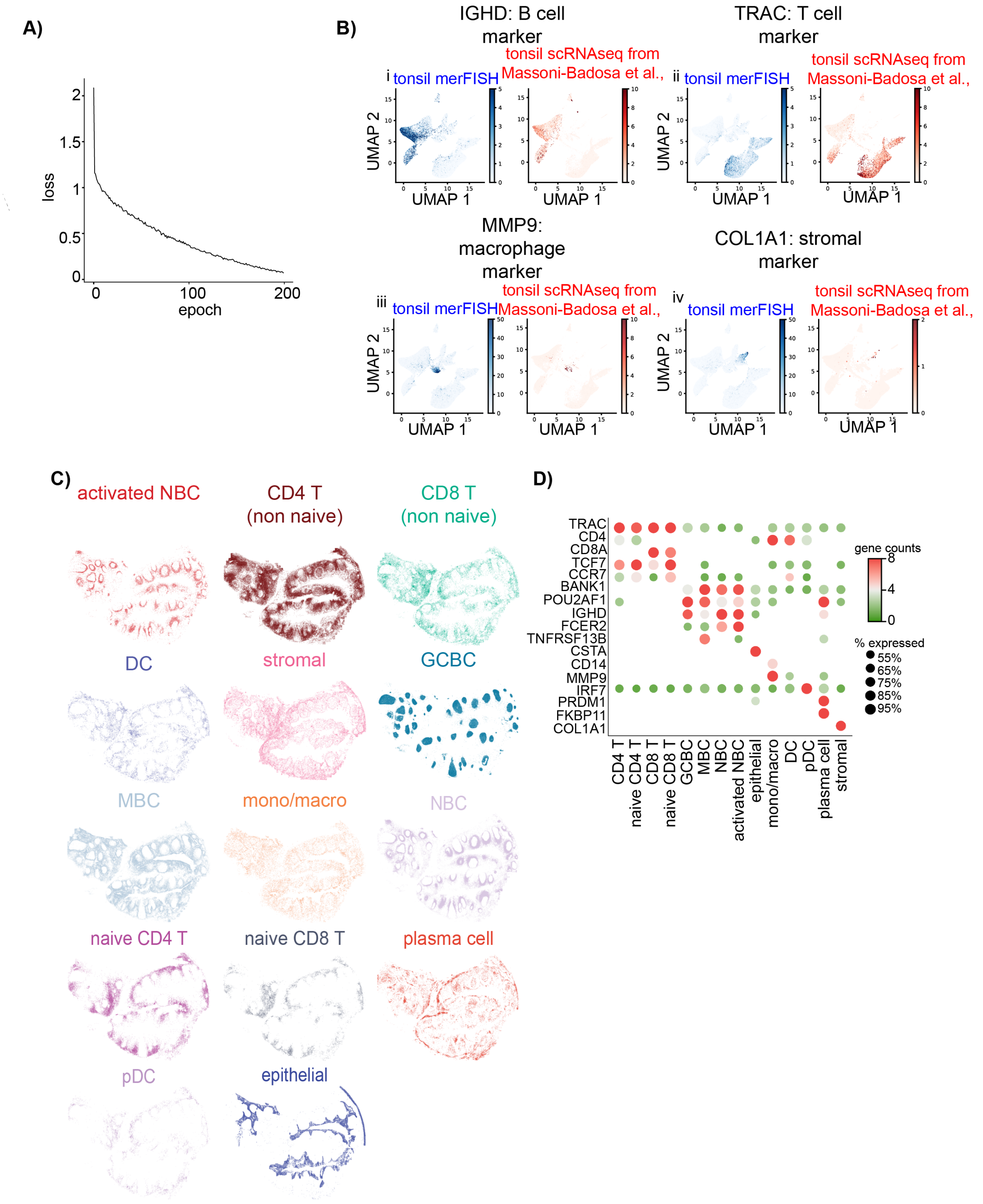

A)
loss
epoch
0
100
200
0.5
1
1.5
2
B)
IGHD: B cell marker
TRAC: T cell marker
MMP9: macrophage marker
COL1A1: stromal marker
i
ii
iii
iv
tonsil merFISH
tonsil scRNAseq from Massoni-Badosa et al.,
UMAP 1
UMAP 2
C)
activated NBC
CD4 T (non naive)
CD8 T (non naive)
DC
stromal
GCBC
MBC
mono/macro
NBC
naive CD4 T
naive CD8 T
plasma cell
pDC
epithelial
D)
TRAC
CD4
CD8A
TCF7
CCR7
BANK1
POU2AF1
IGHD
FCER2
TNFRSF13B
CSTA
CD14
MMP9
IRF7
PRDM1
FKBP11
COL1A1
CD4 T
naive CD4 T
CD8 T
naive CD8 T
GCBC
MBC
NBC
activated NBC
epithelial
mono/macro
DC
pDC
plasma cell
stromal
gene counts
8
4
0
% expressed
55%
65%
75%
85%
95%

Figure 1: **Co-embedding of tonsil sc-RNAseq and Vizgen merFISH data in the same latent space.** **A**: A typical training loss curve for SPARROW-VAE as a function of 200 training epochs. **B**: Expression of known cell type markers shows comparable patterns between ST and sc-RNAseq in the latent space showing successful co-embedding by cell types. Cells in UMAP are visualized using a blue (ST) or red (scRNA-seq) color gradient that is correlated with their expression of cell type markers: IGHD, a B cell marker (i), TRAC, a T cell marker (ii), MMP9, a macrophage marker (iii) and COL1A1 (iv) a stromal marker. Note the scant presence of COL1A1 positive cells in scRNAseq data, which is expected as structural cells are typically under-represented in scRNA-seq assays. **C**: Individual localization of SPARROW-VAE predicted cell types in the tonsil section EXP429. **D**: Known marker genes show expected enrichment in their respective cell types. Mean transcript counts are mapped onto a color gradient, where green and red indicate low and high expression respectively. Dot size is proportional to the percentage of cells expressing the corresponding gene within the indicated cell type. Percentages below 50% are omitted from the plot for brevity.

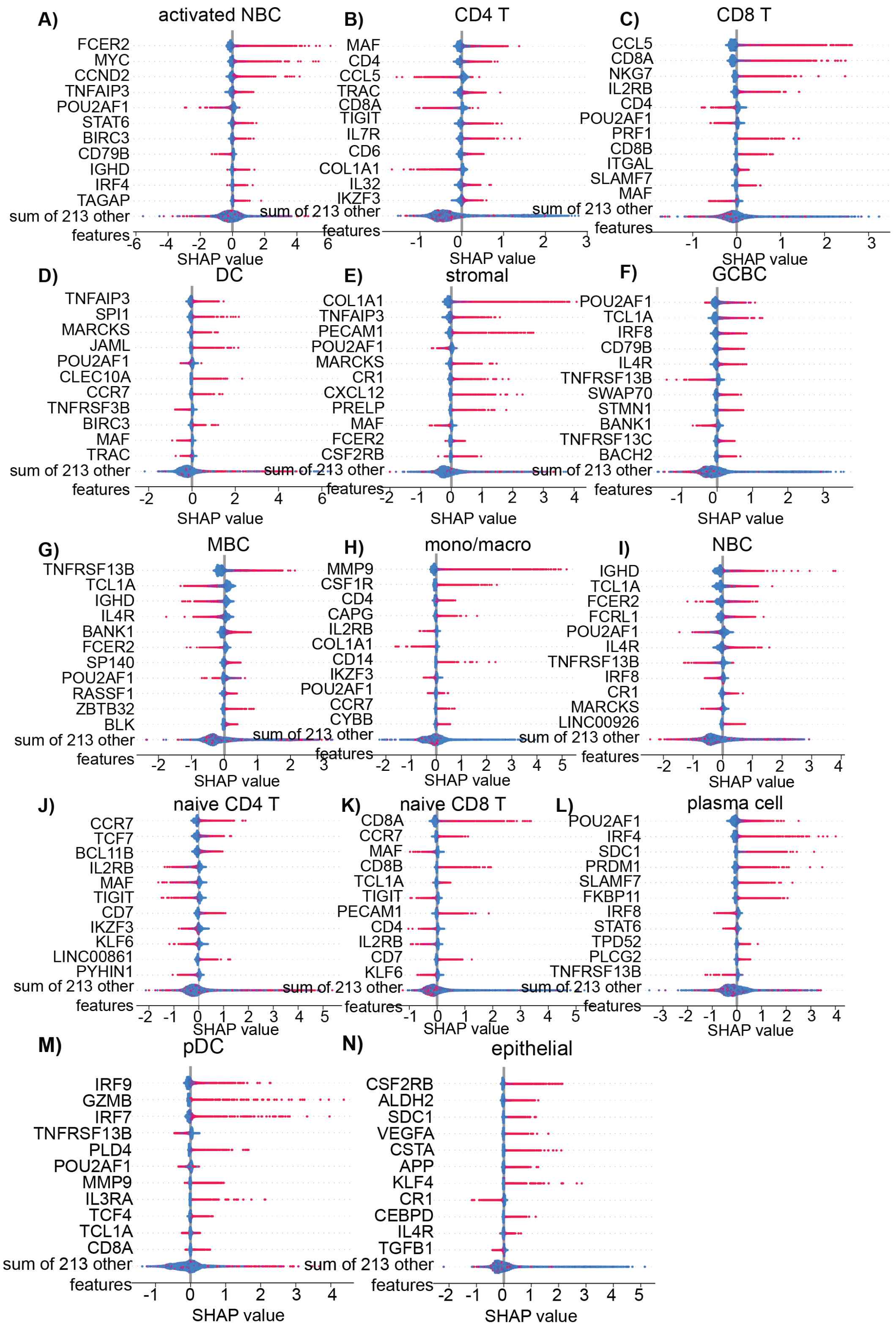


Figure 2: **SHAP scores for top 11 feature genes contributing globally to the SPARROW-VAE cell type inference outcome in tonsil sorted by decreasing importance.**

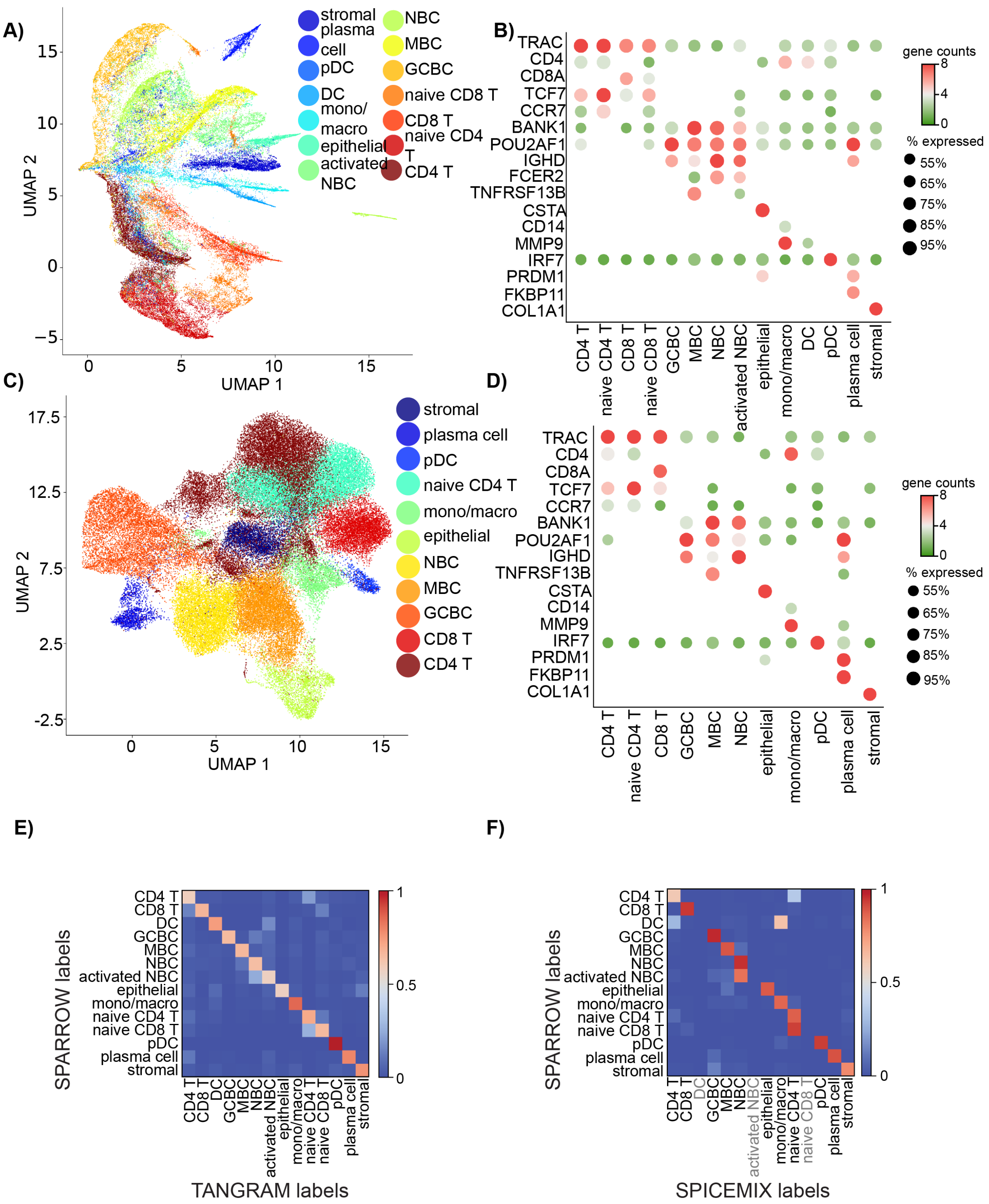

A)
B)
C)
D)
E)
F)
UMAP 1
UMAP 2
stromal
plasma cell
pDC
DC
mono/macro
epithelial
activated NBC
NBC
MBC
GCBC
naive CD8 T
CD8 T
naive CD4 T
CD4 T
TRAC
CD4
CD8A
TCF7
CCR7
BANK1
POU2AF1
IGHD
FCER2
TNFRSF13B
CSTA
CD14
MMP9
IRF7
PRDM1
FKBP11
COL1A1
gene counts
% expressed
55%
65%
75%
85%
95%
SPARROW labels
TANGRAM labels
SPICEMIX labels

Figure 3: **Cell type inference by TANGRAM and SPICEMIX A,C**: 2D UMAP visualization of TANGRAM (A) and SPICEMIX (C) results for tonsil EXP429. **B,D**: Dot plots showing marker gene expression in TANGRAM (B) and SPICEMIX (D) inferred cell types. Mean transcript counts are mapped onto a color gradient, where green and red indicate low and high expression respectively. Dot size is proportional to the percentage of cells expressing the corresponding gene within the indicated cell type. Percentages below 50% are omitted from the plot for brevity. **E-F**: Row normalized confusion matrices showing inference comparisons between SPARROW-VAE and TANGRAM (E) and between SPARROW-VAE and SPICEMIX (F). The percentages of concordantly labelled cells are mapped onto a blue-red color gradient from 0 (0%) to 1 (100%). The matrices are row normalized such rows sum to 1. Note that the cell types absent from SPICEMIX results are marked out in gray.

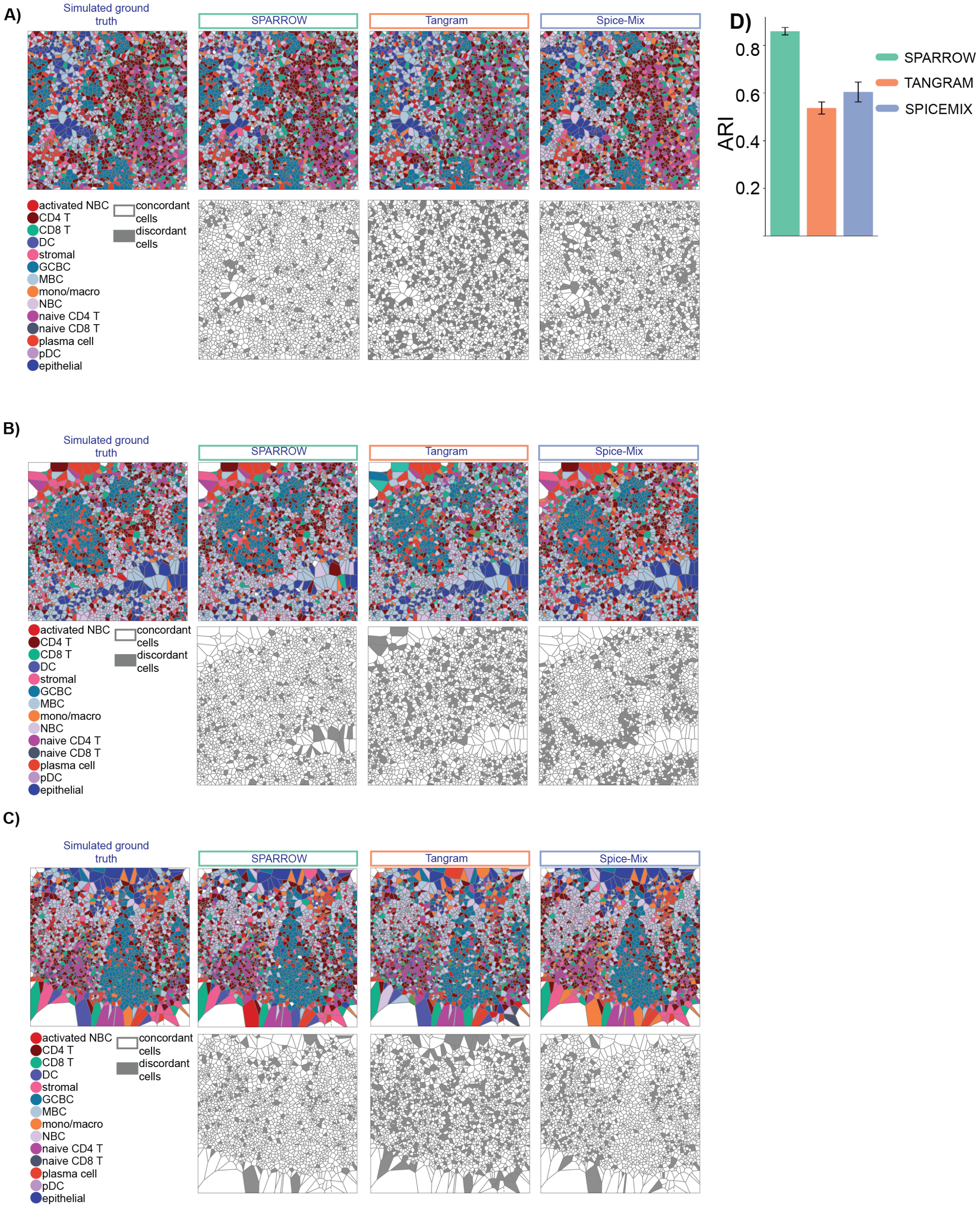

A)
Simulated ground truth
SPARROW
Tangram
Spice-Mix
activated NBC
CD4 T
CD8 T
DC
stromal
GCBC
MBC
mono/macro
NBC
naive CD4 T
naive CD8 T
plasma cell
pDC
epithelial
concordant cells
discordant cells
B)
C)
D)
ARI
0.2
0.4
0.6
0.8
SPARROW
TANGRAM
SPICEMIX

Figure 4: **Evaluation of SPARROW against TANGRAM and SPICEMIX using simulated ground truth cell labels. A-C**: Top row: Voronoi tessellation patterns of the ground truth (leftmost) and inference results from SPARROW-VAE, TANGRAM and SPICEMIX. Bottom row: concordantly (white) and discordantly (gray) labelled cells in SPARROW, TANGRAM and SPICEMIX inference results. **D**: A bar plot showing the average ARI of simulations on three fields of view, measuring the similarity between the predicted cell types from SPARROW, TANGRAM and SPICEMIX to the ground truths. Error bars show standard deviations.

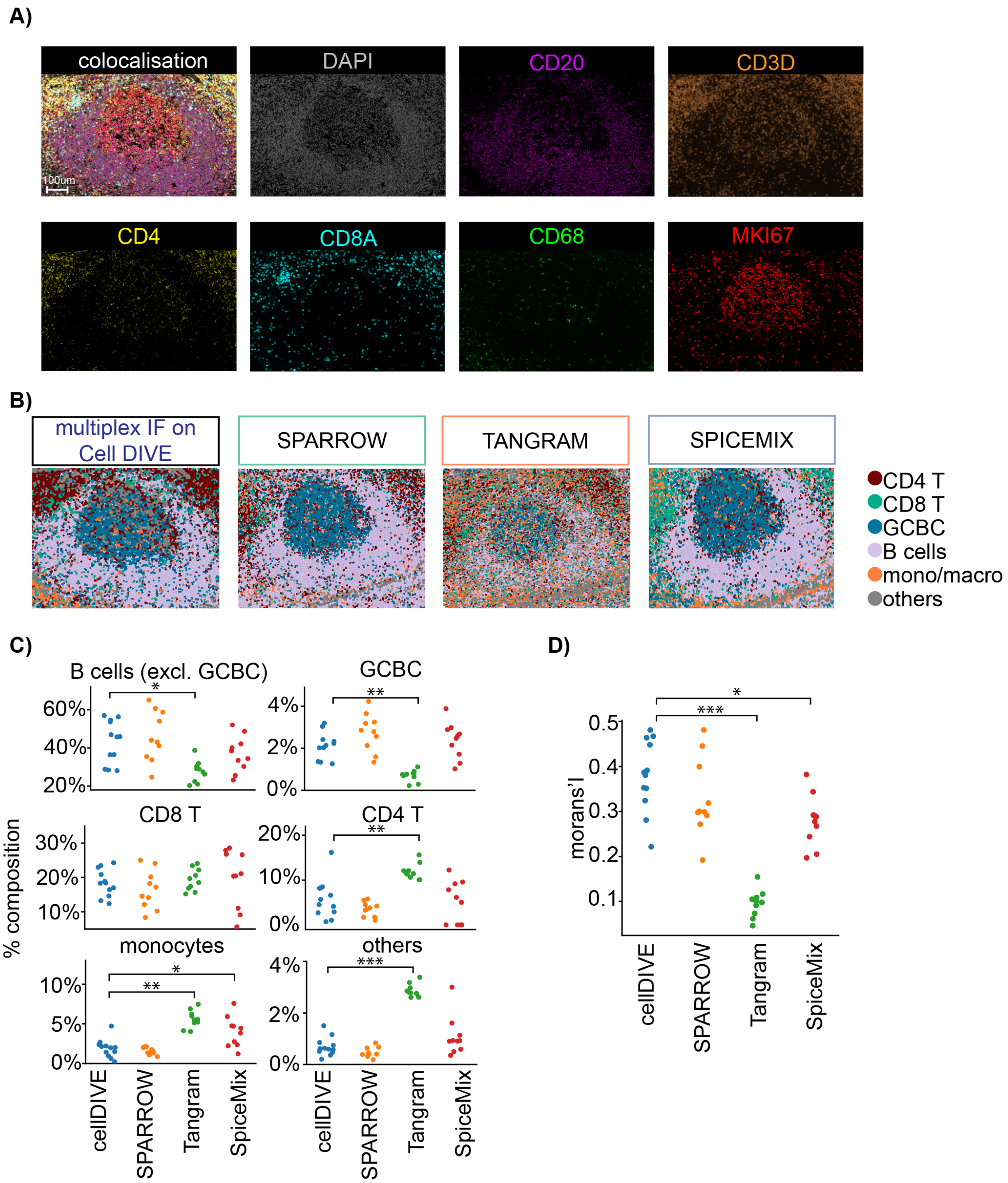
A)
colocalisation
100um
DAPI
CD20
CD3D
CD4
CD8A
CD68
MKI67
B)
multiplex IF on Cell DIVE
SPARROW
TANGRAM
SPICEMIX
CD4 T
CD8 T
GCBC
B cells
mono/macro
others
C)
B cells (excl. GCBC)
GCBC
CD8 T
CD4 T
monocytes
others
% composition
cellDIVE
SPARROW
Tangram
SpiceMix
D)
morans'I
cellDIVE
SPARROW
Tangram
SpiceMix

Figure 5: **Evaluation of SPARROW against TANGRAM and SPICEMIX using orthogonal mxIF imaging of tonsil tissue on Cell DIVE A**: MxIF image of a representative GC in a tonsil tissue section close to that in EXP429 stained with DAPI (white) and immunostained for protein markers: CD20 (purple), CD3D (orange), CD4 (yellow), CD8A (cyan), CD68 (green) and MKI67 (red). **B**: Manual cell typing results in and around a select GC in the Cell DIVE image were shown in a colored coded Voronoi diagram. Cell labels inferred by SPARROW, TANGRAM and SPICEMIX for the same germinal center in EXP429 are shown for visual comparison. **C**: Distributions of percentages of total cells constituted by cell types of interest across 10 GCs inferred from the Cell DIVE image by manual annotation and merFISH image (EXP429) by SPARROW-VAE, TANGRAM and SPICEMIX. **D**: Distributions of Moran's I showing spatial autocorrelation of called cell types in Cell DIVE, SPARROW-VAE, TANGRAM and SPICEMIX results. For **C** and **D** p values were calculated from Mann-Whitney U tests. * $p < 5 \times 10^{-2}$ **$p < 5 \times 10^{-3}$ ***$p < 5 \times 10^{-4}$.

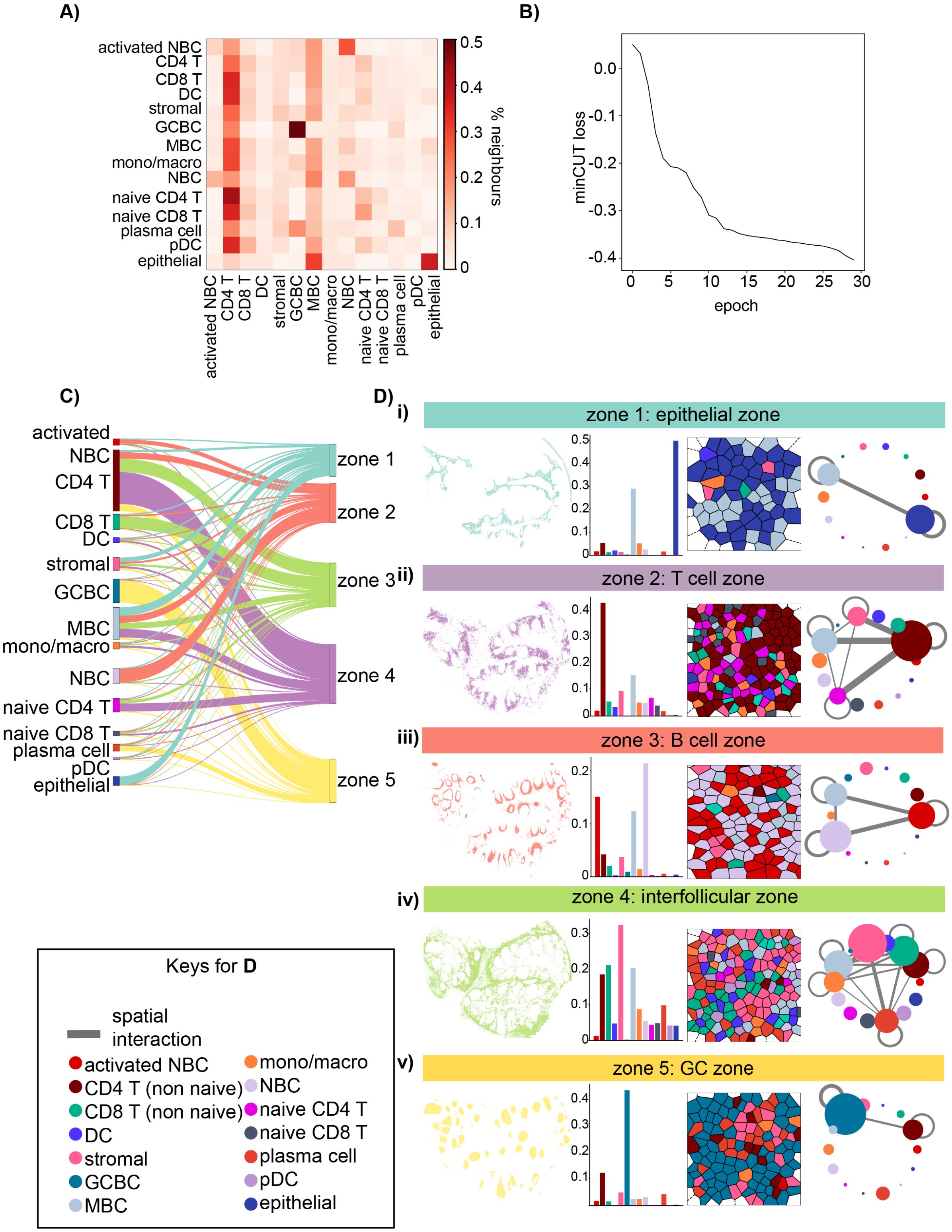
A)
B)
C)
D)
% neighbours
minCUT loss
epoch
activated NBC
CD4 T
CD8 T
DC
stromal
GCBC
MBC
mono/macro
NBC
naive CD4 T
naive CD8 T
plasma cell
pDC
epithelial
zone 1
zone 2
zone 3
zone 4
zone 5
i)
zone 1: epithelial zone
ii)
zone 2: T cell zone
iii)
zone 3: B cell zone
iv)
zone 4: interfollicular zone
v)
zone 5: GC zone
Keys for D
spatial interaction
activated NBC
CD4 T (non naive)
CD8 T (non naive)
DC
stromal
GCBC
MBC
mono/macro
NBC
naive CD4 T
naive CD8 T
plasma cell
pDC
epithelial

Figure 6: **SPARROW identifies microenvironment zones representing distinct cellular neighborhoods.** **A**: A heatmap showing the tissue-wide neighborhood constitution for each cell type. The heatmap is row normalized so that the rows sum to 100%. The pixels represent the neighborhood composition by cell types for the cell types indicated as row names. For instance, the pixel at the intersection between the row for activated NBC and the column for NBC represents the percentage constituted by NBCs among all neighbors of activated NBCs. **B**: A typical training loss curve for SPARROW-GAT as a function of 30 training epochs. **C**: A Sankey Diagram showing the cell type composition of five microenvironment zones. Node height is proportional to the abundance of the represented category. Connector thickness is proportional to the percentage of each cell type present within the respective zone. **D**: Left to right: spatial localization patterns of cells assigned to microenvironment zones 1-5 (i-v), bar plots showing cell type composition within each microenvironment zone, Voronoi tessellation diagrams of a representative region of the indicated microenvironment zone, where cell boundaries are outlined in black and cell types are color-coded according to the scheme in Figure 1, networks summarizing cellular neighborhoods formed by cell types in individual zones in which nodes represent cell types and edges represent immediate adjacency between connected cell types. Node sizes are proportional to the relative abundance of the corresponding cell type in the zone. Edge widths are proportional to the abundance of the spatial interactions between involved cell types. Edges involving cell types with limited presence in the zone (<10% of total cells in a zone) are omitted for brevity. Zone names are assigned based on prior knowledge on tonsil biology.

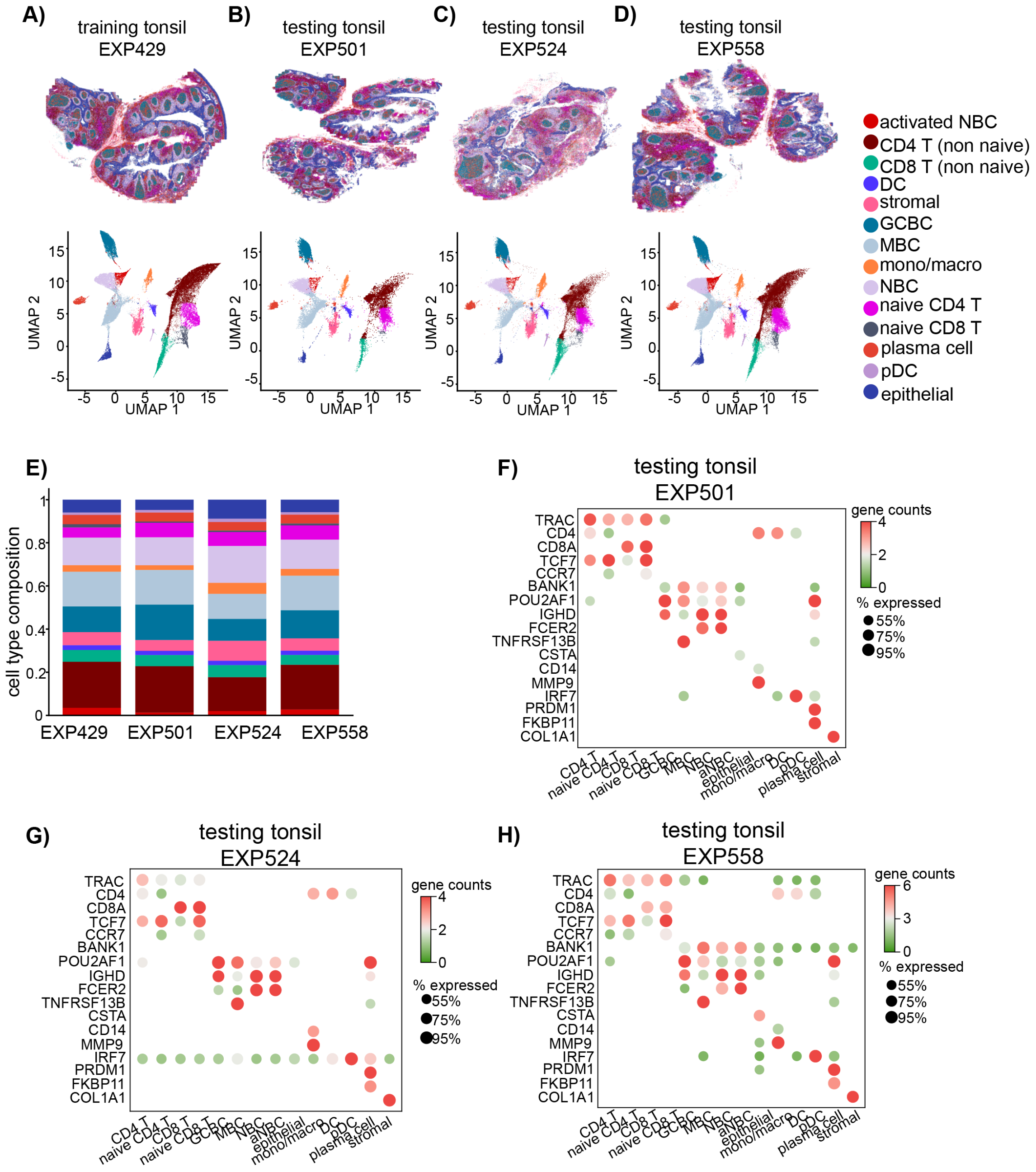


Figure 7: **The common latent space $Z$ constructed by SPARROW-VAE enables co-embedding of multiple sections of the human tonsil A-D**: Cells color-coded according to their cell type prediction in tonsil sections EXP429 (A), EXP501 (B), EXP524 (C) and EXP558 (D) in tissue (top panel) and latent space (bottom panel). The cell label colors are consistent between this figure and Main Figure 1. **E** Stack bar plot showing the cell type composition of tissue sections shown in **A-D**. **F-H**: Dot plots showing marker gene expression in EXP501 (**F**), EXP524 (**G**) and EXP558 (**H**). Mean transcript counts are mapped onto a color gradient, where green and red indicate low and high expression respectively. Dot size is proportional to the percentage of cells expressing the corresponding gene within the indicated cell type. Percentages below 50% are omitted from the plot for brevity.

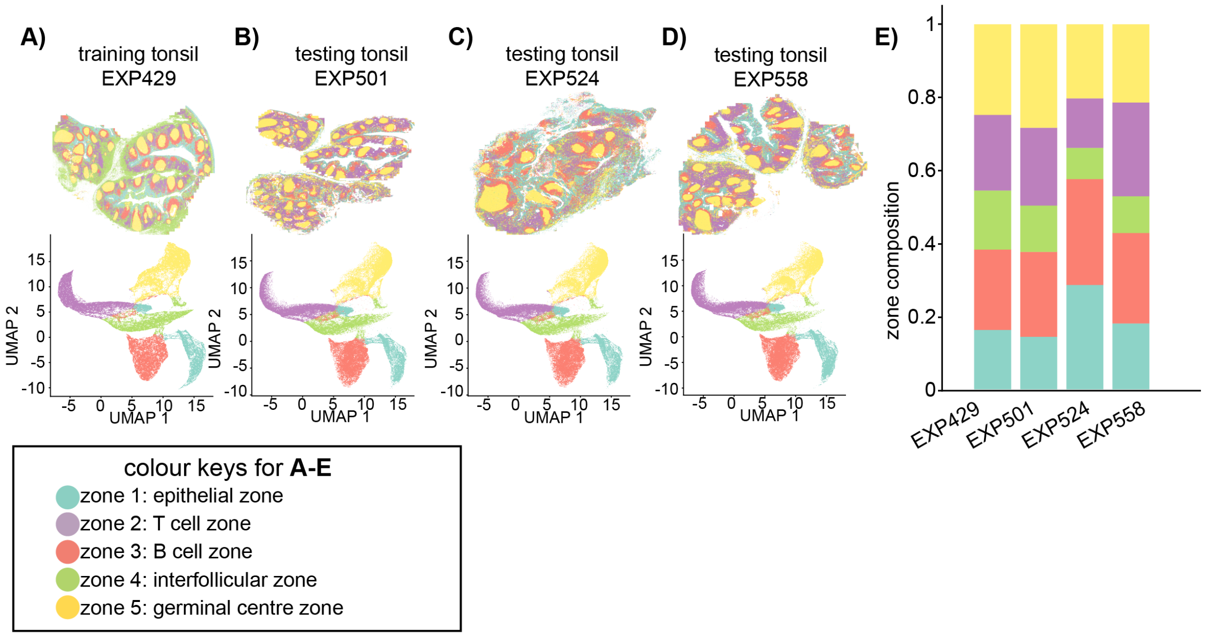


Figure 8: **A-D**: Cells color-coded according to their microenvironment zones prediction in tonsil sections EXP429 (A), EXP501 (B), EXP524 (C) and EXP558 (D) in tissue (top panel) and latent (bottom panel) space. **E**: Stack bar plot showing the microenvironment zone composition of tissue sections shown in **A-D**.

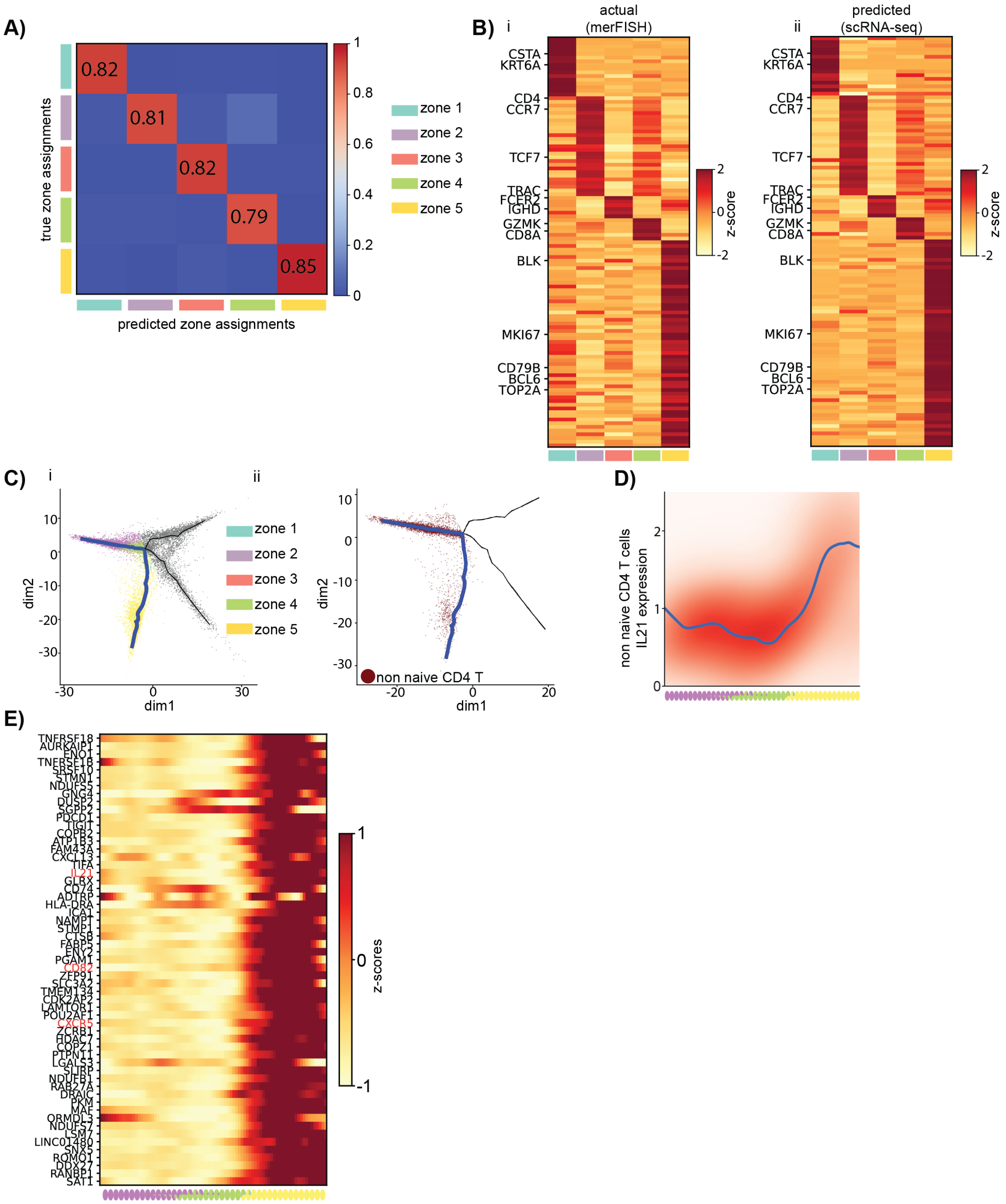
A)
true zone assignments
predicted zone assignments
0.82
0.81
0.82
0.79
0.85
1
0.8
0.6
0.4
0.2
0
zone 1
zone 2
zone 3
zone 4
zone 5
B)
i
actual
(merFISH)
ii
predicted
(scRNA-seq)
CSTA
KRT6A
CD4
CCR7
TCF7
TRAC
FCER2
IGHD
GZMK
CD8A
BLK
MKI67
CD79B
BCL6
TOP2A
z-score
2
0
-2
C)
i
ii
dim1
dim2
non naive CD4 T
D)
non naive CD4 T cells
IL21 expression
E)
TNFRSF18
AURKAIP1
ENO1
TNFRSF1B
SRSF10
STMN1
NDUFS5
GNG4
DUSP2
SGPP2
PDCD1
TIGIT
COPB2
ATP1B3
FAM43A
CXCL13
TIFA
IL21
GLRX
CD74
ADTRP
HLA-DRA
ICA1
NAMPT
STMP1
CTSB
FABP5
ENY2
PGAM1
CD82
ZFP91
SLC3A2
TMEM134
CDK2AP2
LAMTOR1
POU2AF1
CXCR5
ZCRB1
HDAC7
COPZ1
PTPN11
LGALS3
SLIRP
NDUFB1
RAB27A
DRAIC
PKM
MAF
ORMDL3
NDUFS7
LSM7
LINC01480
SNX5
ROMO1
DDX27
RANBP1
SAT1
z-scores
1
0
-1

Figure 9: **Microenvironment zone predictions reveal dynamic transcriptomic changes in cell types across the tissue space.** **A**: A confusion matrix showing the accuracy of microenvironment zone predictions. Matrix values represent row percentages of true vs. predicted zone comparisons. True zone assignments are microenvironment zone assignments $H$ for the test tonsil tissue (EXP501) inferred using both the expression matrix and spatial adjacency, while predicted zone assignments are $H'$ using only the expression matrix. **B**: Heatmaps of gene expression z-scores for genes with significant zone specific expression patterns in tonsil merFISH data of EXP429 (left) and in predicted zone assignments from tonsil scRNA-seq (right). Zone specificity is established through the Kruskal-Wallis test ($p < 0.05$), followed by Dunn's test with Bonferroni correction to determine the zone in which the gene is enriched. Each row represents a marker gene. Representative zone specific marker genes are marked out on the y-axis. **C**: i: A principal graph was constructed for SPARROW-GAT latent embedding $H$ of cells in tonsil EXP409. Cells in the graph space are represented by dots color-coded according to their respective microenvironment zone assignment. The select trajectory shown on the x-axes of Main Figure 1 F and G is shown in blue. ii: Non naive CD4 T cells mapped on the select trajectory are plotted as dark red dots, which show a bimodal distribution characterised by a predominant presence in zone 2, the T cell zone, and zone 4, the interfollicular zone, with a reduced presence in zone 5, the GC zone. **D**: IL21 expression of non naive CD4 T cells in scRNA-seq along the select trajectory (dark blue in **C**). Below the x-axis, pie charts display the percentage of cells with color-coded zone predictions projected to the select trajectory in the graph. **E**: A heatmap of gene expression z-scores in non naive CD4 T cells along the selected trajectory (dark blue in **C**) for genes with significant expression increases in zone 5, the GC zone, as determined by the Kruskal-Wallis test followed by Dunn's test with Bonferroni correction ($p < 0.05$). Genes that are known to be important for non naive CD4 T cell migration into GCs are marked out in red.

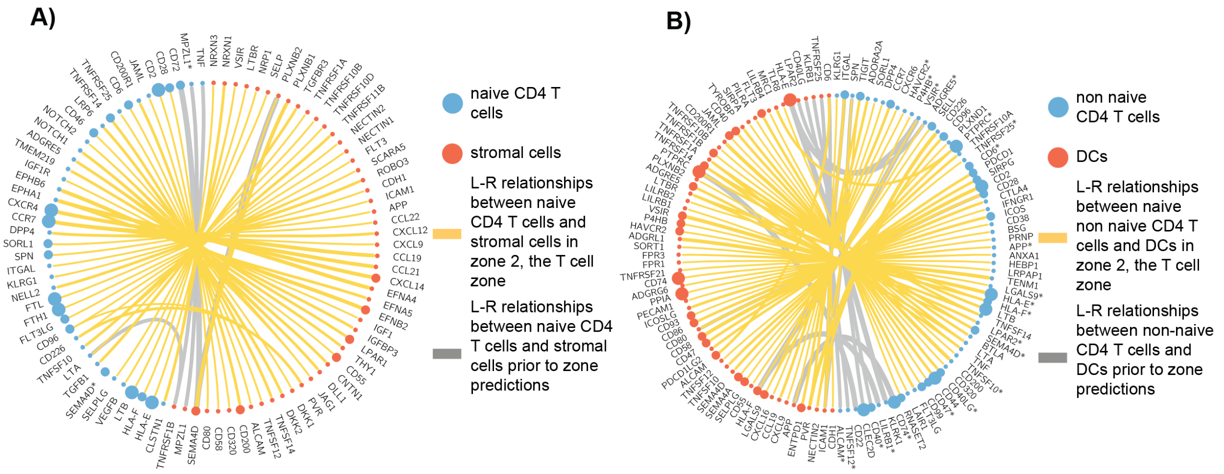


Figure 10: **Biologically significant L-R relationships between specific cell types are elucidated upon microenvironment zone predictions.** Circos plots showing significant L-R pairs between naive CD4 T cells and stromal cells (**A**) and between non naive CD4 T cells and DCs (**B**) in zone 2, the T cell zone. Pairs were identified using cellphonedb modelling. Significance was fined as p values $< 0.05$. The widths of connectors are positively correlated with interaction scores. In cases where the same genes are expressed in both cell types, the genes associated with cell types marked out by red nodes are marked with asterisks.

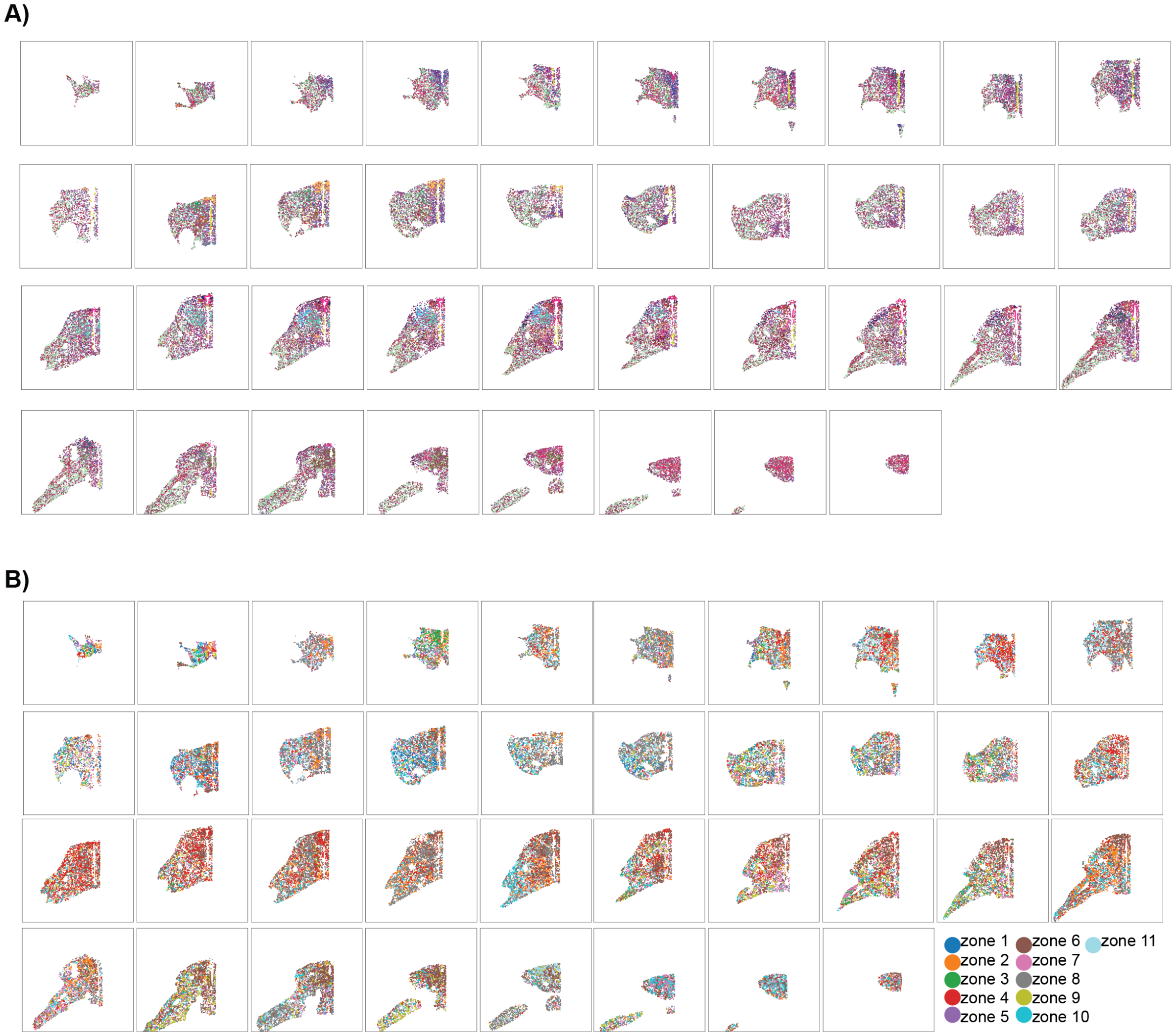


Figure 11: **Spatial visualisation of cells in sequential slices of the mouse hypothalamus in Zhang et al., 2023, colour coded by cell type subclasses as defined in the original publication (A) and SPARROW microenvironment zones (B).** Colour keys used in **A** are consistent with the original publication.

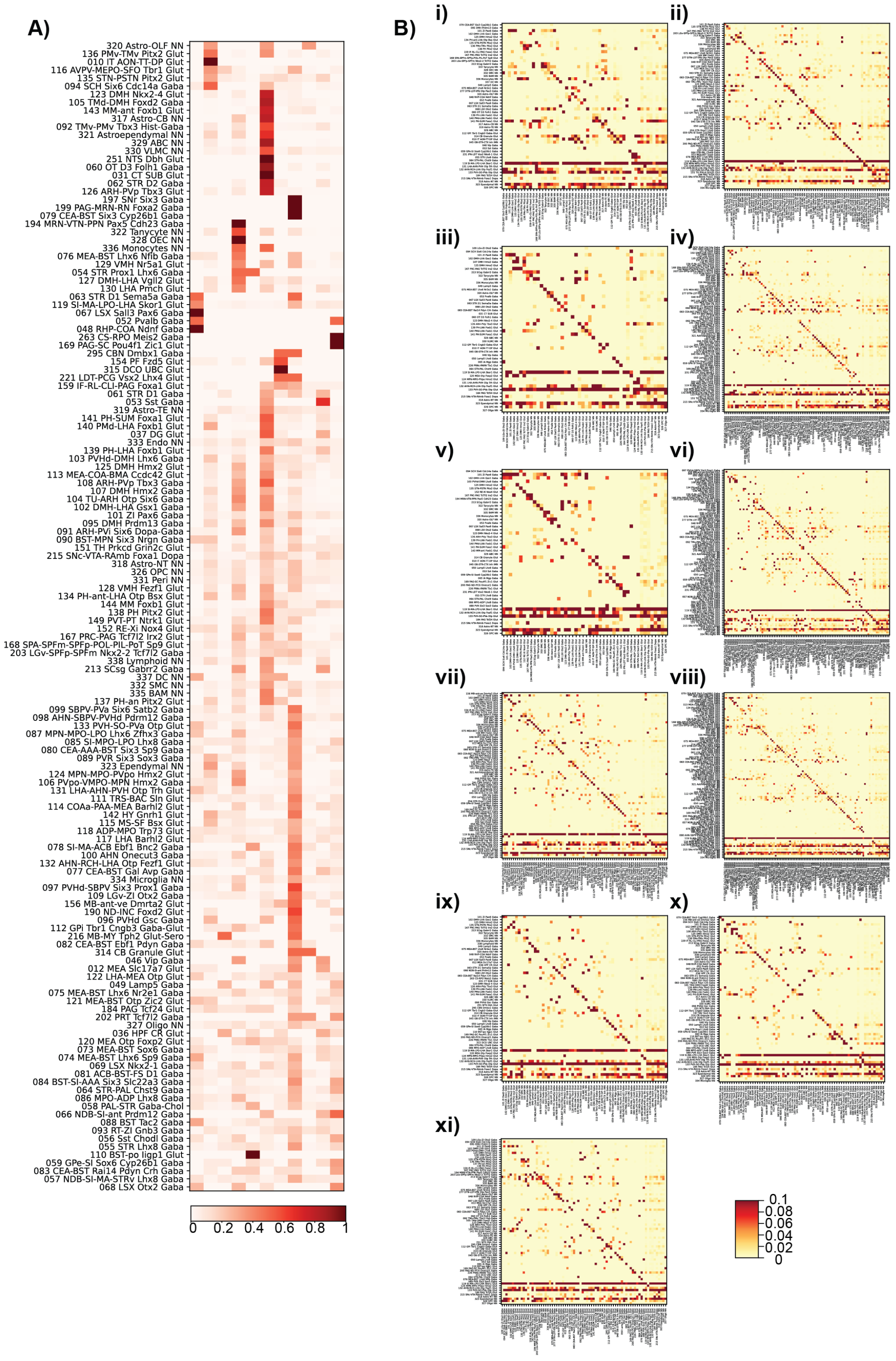
A)
320 Astro-OLF NN
136 PMv-TMv Pitx2 Glut
010 IT AON-TT-DP Glut
116 AVPV-MEPO-SFO Tbr1 Glut
135 STN-PSTN Pitx2 Glut
094 SCH Six6 Cdc14a Gaba
123 DMH Nkx2-4 Glut
105 TMd-DMH Foxd2 Gaba
143 MM-ant Foxb1 Glut
317 Astro-CB NN
092 TMv-PMv Tbx3 Hist-Gaba
321 Astroependymal NN
329 ABC NN
330 VLMC NN
251 NTS Dbh Glut
060 OT D3 Folh1 Gaba
031 CT SUB Glut
062 STR D2 Gaba
126 ARH-PVp Tbx3 Glut
197 SNr Six3 Gaba
199 PAG-MRN-RN Foxa2 Gaba
079 CEA-BST Six3 Cyp26b1 Gaba
194 MRN-VTN-PPN Pax5 Cdh23 Gaba
322 Tanycyte NN
328 OEC NN
336 Monocytes NN
076 MEA-BST Lhx6 Nfib Gaba
129 VMH Nr5a1 Glut
054 STR Prox1 Lhx6 Gaba
127 DMH-LHA Vgll2 Glut
130 LHA Pmch Glut
063 STR D1 Sema5a Gaba
119 SI-MA-LPO-LHA Skor1 Glut
067 LSX Sall3 Pax6 Gaba
052 Pvalb Gaba
048 RHP-COA Ndnf Gaba
263 CS-RPO Meis2 Gaba
169 PAG-SC Pou4f1 Zic1 Glut
295 CBN Dmbx1 Gaba
154 PF Fzd5 Glut
315 DCO UBC Glut
221 LDT-PCG Vsx2 Lhx4 Glut
159 IF-RL-CLI-PAG Foxa1 Glut
061 STR D1 Gaba
053 Sst Gaba
319 Astro-TE NN
141 PH-SUM Foxa1 Glut
140 PMd-LHA Foxb1 Glut
037 DG Glut
333 Endo NN
139 PH-LHA Foxb1 Glut
103 PVHd-DMH Lhx6 Gaba
125 DMH Hmx2 Glut
113 MEA-COA-BMA Ccdc42 Glut
108 ARH-PVp Tbx3 Gaba
107 DMH Hmx2 Gaba
104 TU-ARH Otp Six6 Gaba
102 DMH-LHA Gsx1 Gaba
101 ZI Pax6 Gaba
095 DMH Prdm13 Gaba
091 ARH-PVi Six6 Dopa-Gaba
090 BST-MPN Six3 Nrgn Gaba
151 TH Prkcd Grin2c Glut
215 SNc-VTA-RAmb Foxa1 Dopa
318 Astro-NT NN
326 OPC NN
331 Peri NN
128 VMH Fezf1 Glut
134 PH-ant-LHA Otp Bsx Glut
144 MM Foxb1 Glut
138 PH Pitx2 Glut
149 PVT-PT Ntrk1 Glut
152 RE-Xi Nox4 Glut
167 PRC-PAG Tcf7l2 Irx2 Glut
168 SPA-SPFm-SPFp-POL-PIL-PoT Sp9 Glut
203 LGv-SPFp-SPFm Nkx2-2 Tcf7l2 Gaba
338 Lymphoid NN
213 SCsg Gabrr2 Gaba
337 DC NN
332 SMC NN
335 BAM NN
137 PH-an Pitx2 Glut
099 SBPV-PVa Six6 Satb2 Gaba
098 AHN-SBPV-PVHd Pdrm12 Gaba
133 PVH-SO-PVa Otp Glut
087 MPN-MPO-LPO Lhx6 Zfhx3 Gaba
085 SI-MPO-LPO Lhx8 Gaba
080 CEA-AAA-BST Six3 Sp9 Gaba
089 PVR Six3 Sox3 Gaba
323 Ependymal NN
124 MPN-MPO-PVpo Hmx2 Glut
106 PVpo-VMPO-MPN Hmx2 Gaba
131 LHA-AHN-PVH Otp Trh Glut
111 TRS-BAC Sln Glut
114 COAa-PAA-MEA Barhl2 Glut
142 HY Gnrh1 Glut
115 MS-SF Bsx Glut
118 ADP-MPO Trp73 Glut
117 LHA Barhl2 Glut
078 SI-MA-ACB Ebf1 Bnc2 Gaba
100 AHN Onecut3 Gaba
132 AHN-RCH-LHA Otp Fezf1 Glut
077 CEA-BST Gal Avp Gaba
334 Microglia NN
097 PVHd-SBPV Six3 Prox1 Gaba
109 LGv-ZI Otx2 Gaba
156 MB-ant-ve Dmrta2 Glut
190 ND-INC Foxd2 Glut
096 PVHd Gsc Gaba
112 GPi Tbr1 Cngb3 Gaba-Glut
216 MB-MY Tph2 Glut-Sero
082 CEA-BST Ebf1 Pdyn Gaba
314 CB Granule Glut
046 Vip Gaba
012 MEA Slc17a7 Glut
122 LHA-MEA Otp Glut
049 Lamp5 Gaba
075 MEA-BST Lhx6 Nr2e1 Gaba
121 MEA-BST Otp Zic2 Glut
184 PAG Tcf24 Glut
202 PRT Tcf7l2 Gaba
327 Oligo NN
036 HPF CR Glut
120 MEA Otp Foxp2 Glut
073 MEA-BST Sox6 Gaba
074 MEA-BST Lhx6 Sp9 Gaba
069 LSX Nkx2-1 Gaba
081 ACB-BST-FS D1 Gaba
084 BST-SI-AAA Six3 Slc22a3 Gaba
064 STR-PAL Chst9 Gaba
086 MPO-ADP Lhx8 Gaba
058 PAL-STR Gaba-Chol
066 NDB-SI-ant Prdm12 Gaba
088 BST Tac2 Gaba
093 RT-ZI Gnb3 Gaba
056 Sst Chodl Gaba
055 STR Lhx8 Gaba
110 BST-po Iigp1 Glut
059 GPe-SI Sox6 Cyp26b1 Gaba
083 CEA-BST Rai14 Pdyn Crh Gaba
057 NDB-SI-MA-STRv Lhx8 Gaba
068 LSX Otx2 Gaba
0 0.2 0.4 0.6 0.8 1
B)
i)
ii)
iii)
iv)
v)
vi)
vii)
viii)
ix)
x)
xi)
0.1
0.08
0.06
0.04
0.02
0

Figure 12: **SPARROW-GAT delineated microenvironment zones in the mouse hypothalamus are characterized by unique cell type composition and cell neighborhoods.** **A**: A heatmap showing cell type composition of mouse hypothalamus microenvironment zones. **B**: Heatmaps summarizing neighborhood compositions in zones 1-11. Rows and columns indicate spatially adjacent cell types. Pixels indicate numbers of neighboring cells constituted by the cell types specified.

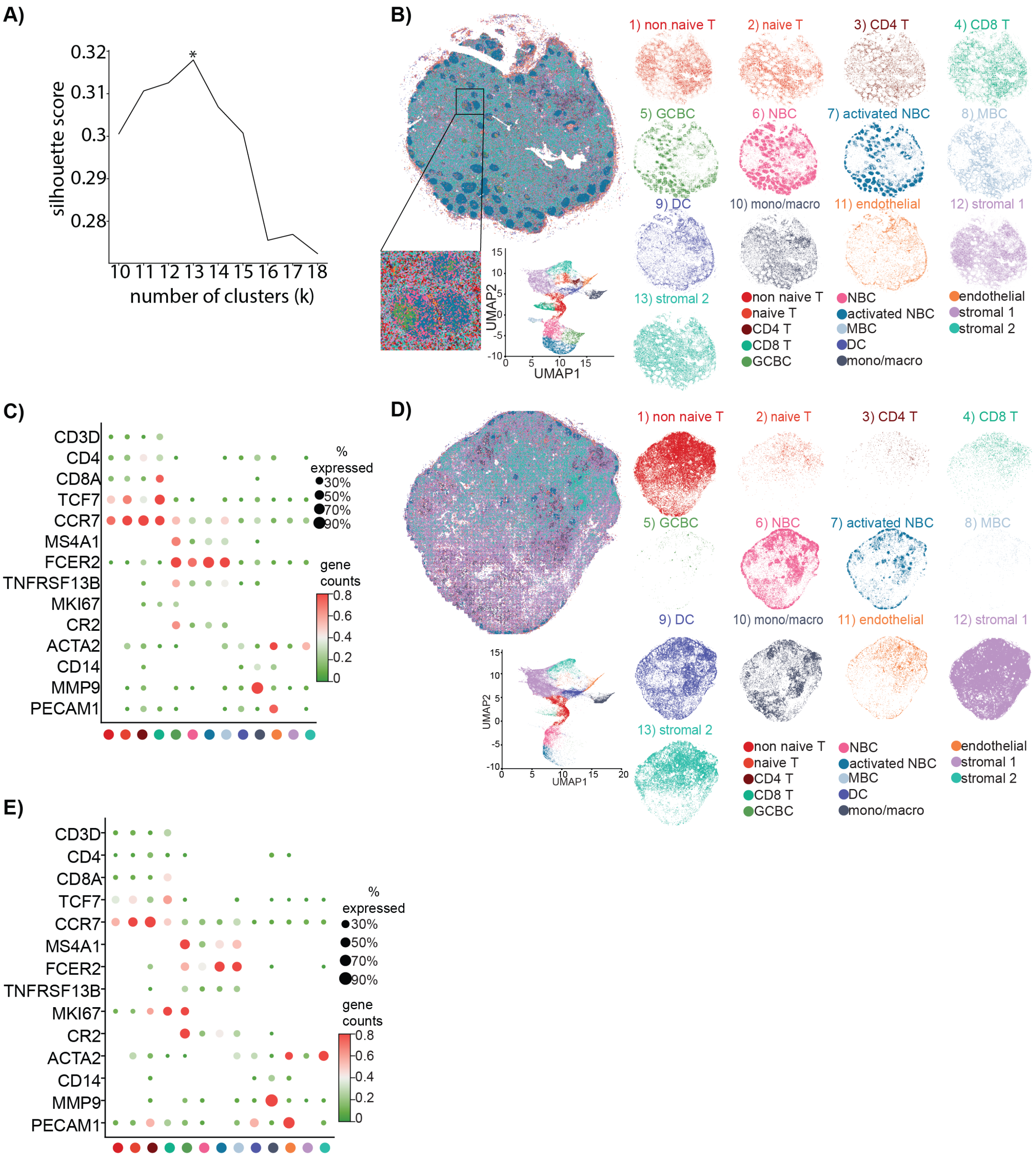
A)
silhouette score
0.32
0.31
0.3
0.29
0.28
*
10 11 12 13 14 15 16 17 18
number of clusters (k)
B)
1) non naive T
2) naive T
3) CD4 T
4) CD8 T
5) GCBC
6) NBC
7) activated NBC
8) MBC
9) DC
10) mono/macro
11) endothelial
12) stromal 1
13) stromal 2
UMAP1
UMAP2
non naive T
naive T
CD4 T
CD8 T
GCBC
NBC
activated NBC
MBC
DC
mono/macro
endothelial
stromal 1
stromal 2
C)
CD3D
CD4
CD8A
TCF7
CCR7
MS4A1
FCER2
TNFRSF13B
MKI67
CR2
ACTA2
CD14
MMP9
PECAM1
% expressed
30%
50%
70%
90%
gene counts
0.8
0.6
0.4
0.2
0
D)
1) non naive T
2) naive T
3) CD4 T
4) CD8 T
5) GCBC
6) NBC
7) activated NBC
8) MBC
9) DC
10) mono/macro
11) endothelial
12) stromal 1
13) stromal 2
UMAP1
UMAP2
non naive T
naive T
CD4 T
CD8 T
GCBC
NBC
activated NBC
MBC
DC
mono/macro
endothelial
stromal 1
stromal 2
E)
CD3D
CD4
CD8A
TCF7
CCR7
MS4A1
FCER2
TNFRSF13B
MKI67
CR2
ACTA2
CD14
MMP9
PECAM1
% expressed
30%
50%
70%
90%
gene counts
0.8
0.6
0.4
0.2
0

Figure 13: **Application of SPARROW-VAE to human LNs A**: Silhouette scores for k-means clustering on SPARROW-VAE latent representation $Z$ of LN EXP621.Optimum cluster number is marked by the asterisk. **B**: SPARROW-VAE inferred cell types in the mediastinal LN tissue section EXP621 are visualized in tissue space collectively and in individual panels. A 1000 pixel $\times$ 1000 pixel window of the tissue section was zoomed in to show details. Latent representation $Z$ of the cells in the selected window is visualized in a 2D UMAP space. **C**: Known marker genes show expected enrichment in their respective cell types. Mean transcript counts are mapped onto a color gradient, where green and red indicate low and high expression respectively. Dot size is proportional to the percentage of cells expressing the corresponding gene within the indicated cell type. Percentages below 10% are omitted from the plot for brevity. **D**: SPARROW-VAE model trained on the mediastinal LN section (EXP621) was applied to the mesenteric LN section (EXP505). Cell Latent representations $Z$ are visualized 2D UMAP plots and color-coded according to their cell type labels. **E**: Marker genes for the respective cell types show expected enrichment in EXP505. Mean transcript counts are mapped onto a color gradient, where green and red indicate low and high expression respectively. Dot size is proportional to the percentage of cells expressing the corresponding gene within the indicated cell type. Percentages below 10% are omitted from the plot for brevity.

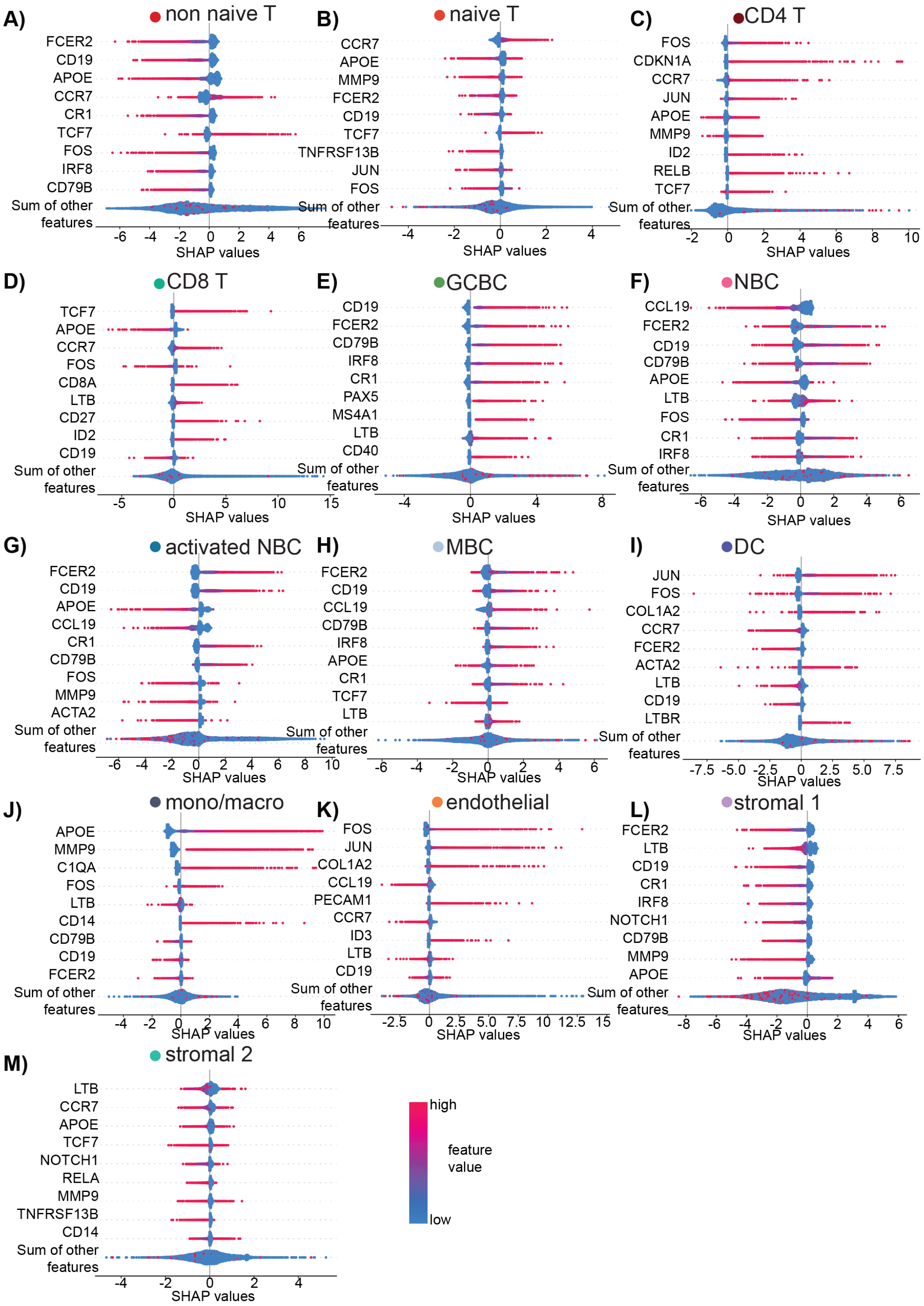


Figure 14: **SHAP scores for top 9 feature genes contributing globally to the SPARROW-VAE cell type inference outcome in the LN EXP621 sorted by decreasing importance.**

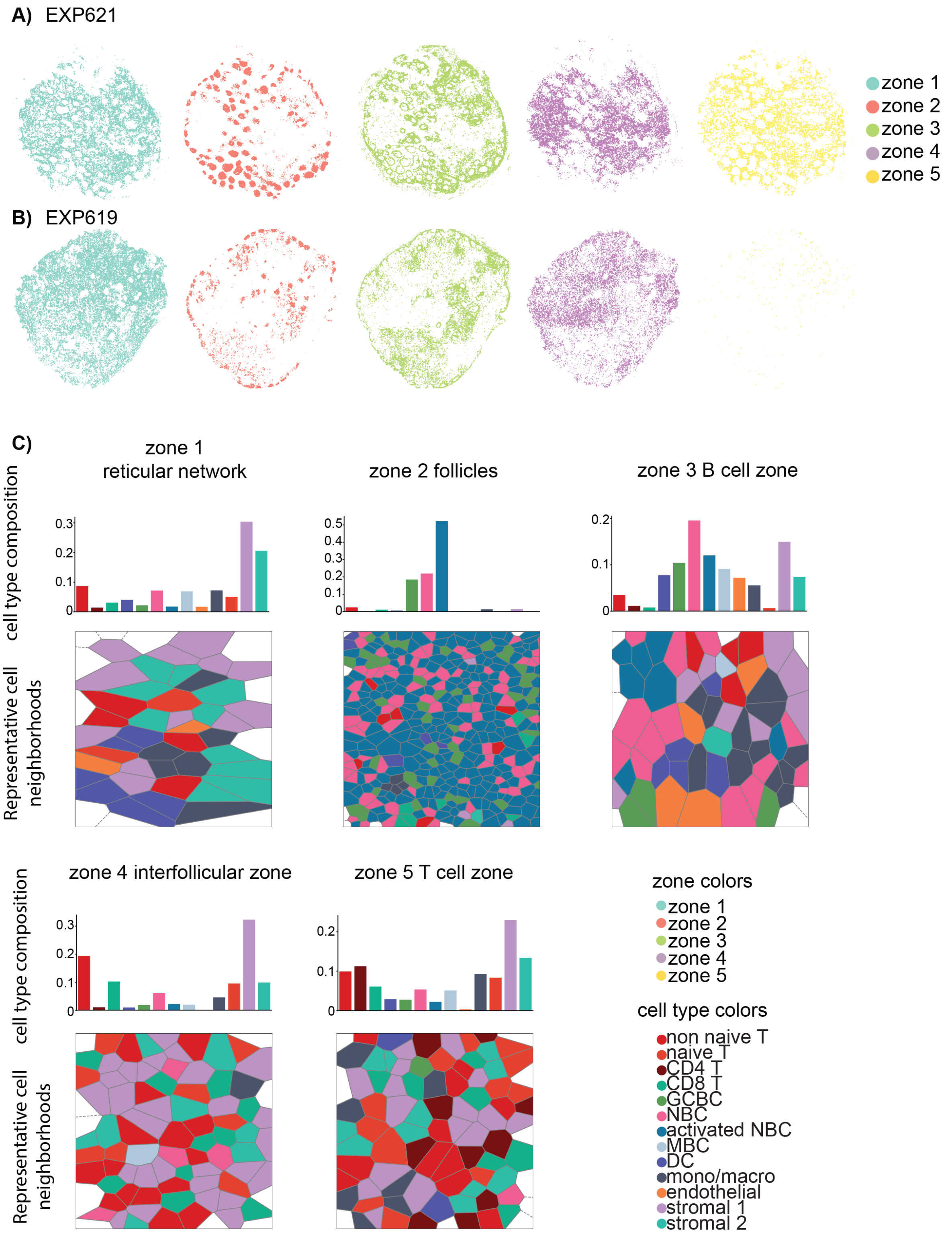


Figure 15: **Application of SPARROW-GAT to human LNs** Spatial localization of microenvironment zones viewed individually in the mediastinal LN (EXP621) and mesenteric LN (EXP505) (**A** and **B**). **C**: Cell type composition of individual microenvironment zones in the LN EXP621. Top row: bar plots showing cell type composition of zones 1-5. Bottom row: Representative Voronoi diagrams where gray lines mark cell boundaries and cells are color coded based on their cell type. Areas were deliberately selected to consist of single zones.

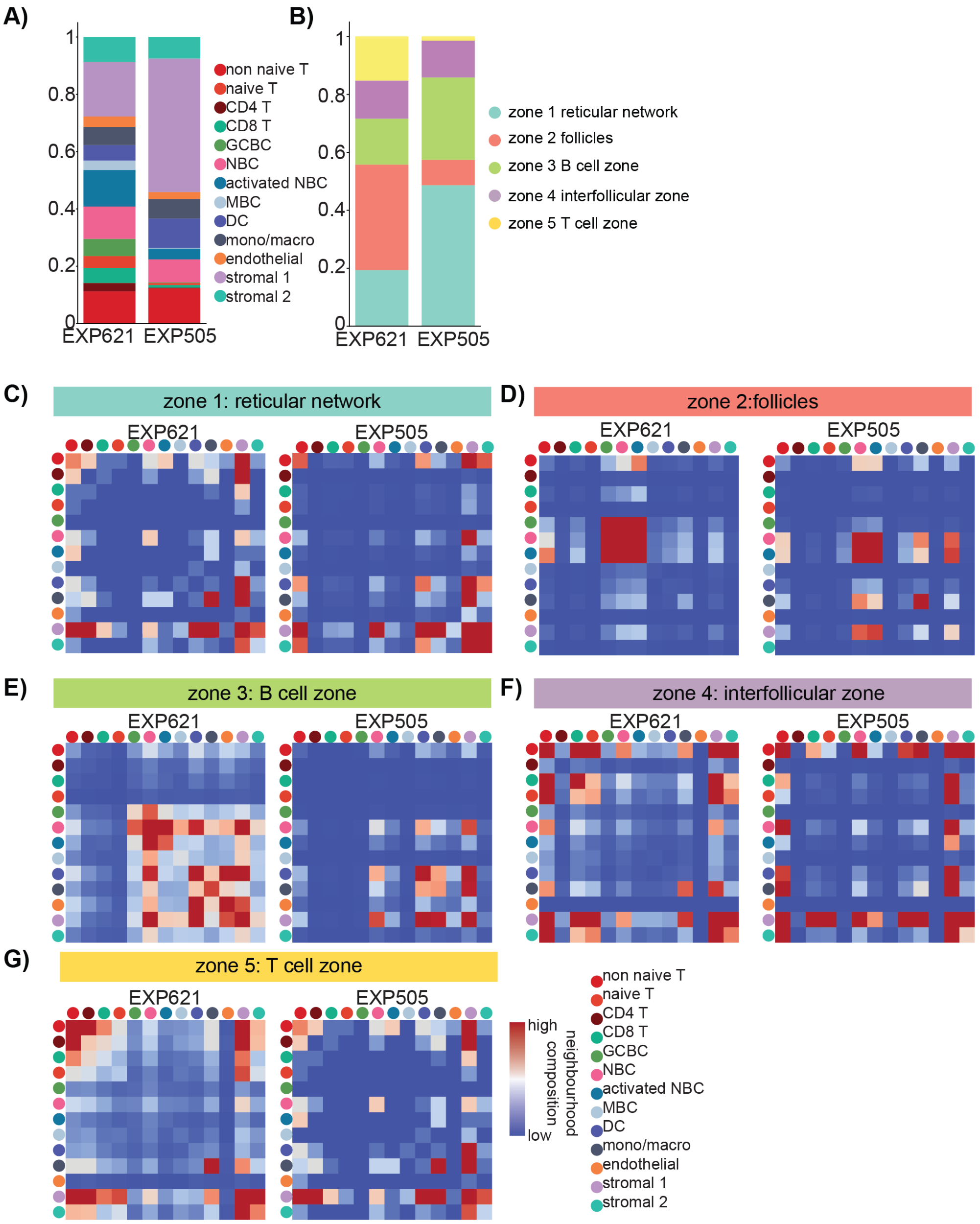


Figure 16: **SPARROW facilitates direct comparisons of cell type composition and microenvironment zones between two human LNs.** Bar plots showing cell type (**A**) and microenvironment zone (**B**) composition of EXP621 and EXP505. **C-G**: Heatmaps summarizing neighborhood compositions in zones 1-5 in EXP621 and EXP505 show overall similar interaction patterns between the two LN of distinct anatomical origins. Rows and columns indicate spatially adjacent cell types. Pixels indicate numbers of neighboring cells involving cell types specified.

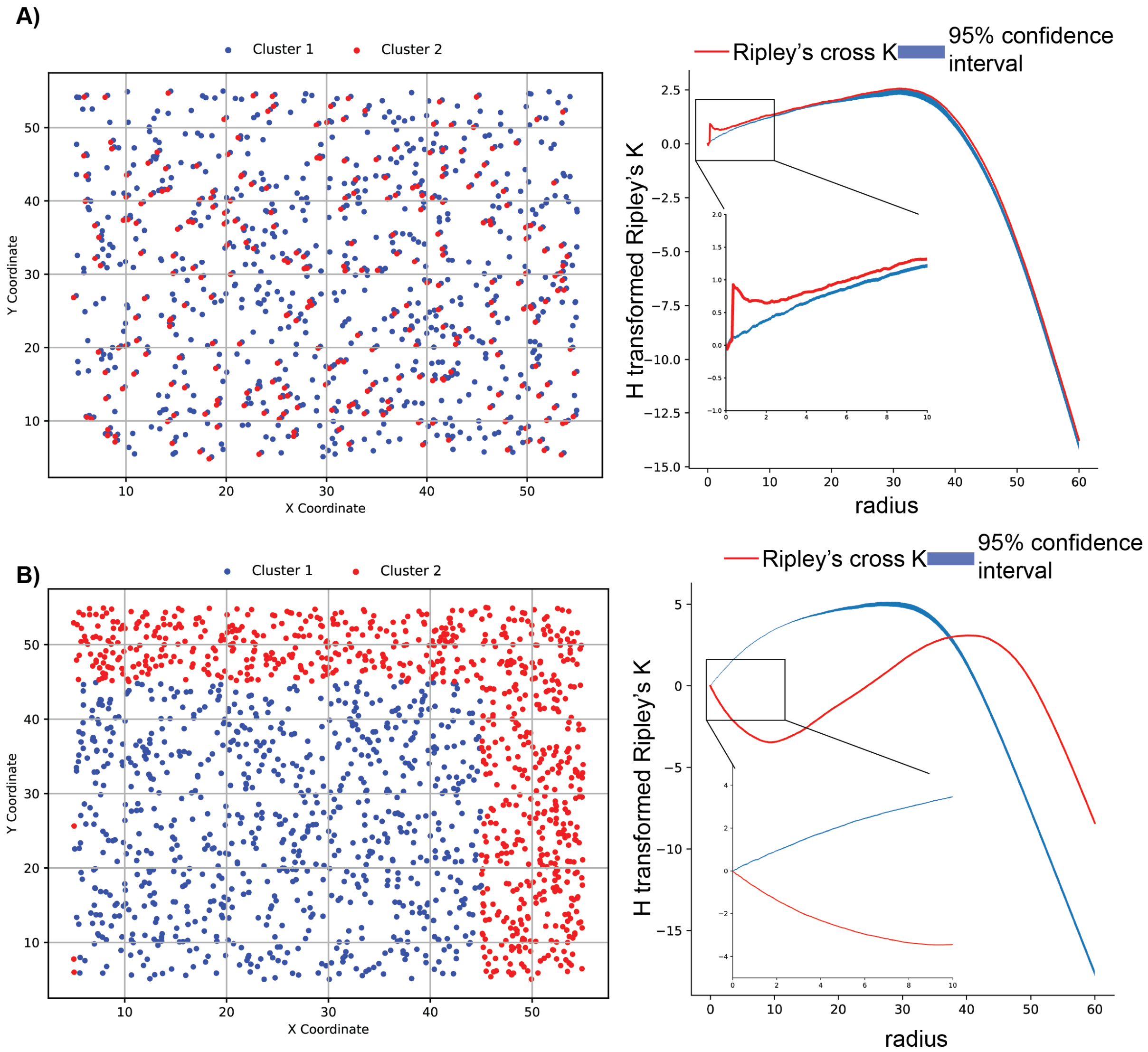


Figure 17: **Simulated spatially co-localised or exclusive point patterns and their Ripley's cross K curves.** Two types of point patterns were generated in **A** and **B**. In **A**, each red cell is accompanied by a blue cell one cell radius away, mimicking cell-to-cell interaction. In **B**, red cells and blue cells are spatially exclusive. H-transformed Ripley's cross K of blue cells relative to red cells as a function of increasing distance was computed (red line). 1000 iterations of spatial random patterns were generated by randomly shuffling blue and red labels and H-transformed Ripley's cross K was calculated for every random pattern iteration. The 95% confidence envelope of the random patterns was plotted (blue belt) and compared against the experimental curve (red line). For the point pattern in **A**, Ripley's cross K for the experimental curve is numerically larger than that of the random pattern at the distance of 1 cell radius (supported by the fact that the experimental curve is elevated above the confidence belt at the corresponding distance), indicating statistically significant co-localisation. For the point pattern in **B**, Ripley's cross K for the experimental curve is numerically smaller than that of the random pattern at a range of short distances, indicating statistically significant spatial exclusivity at these distances.

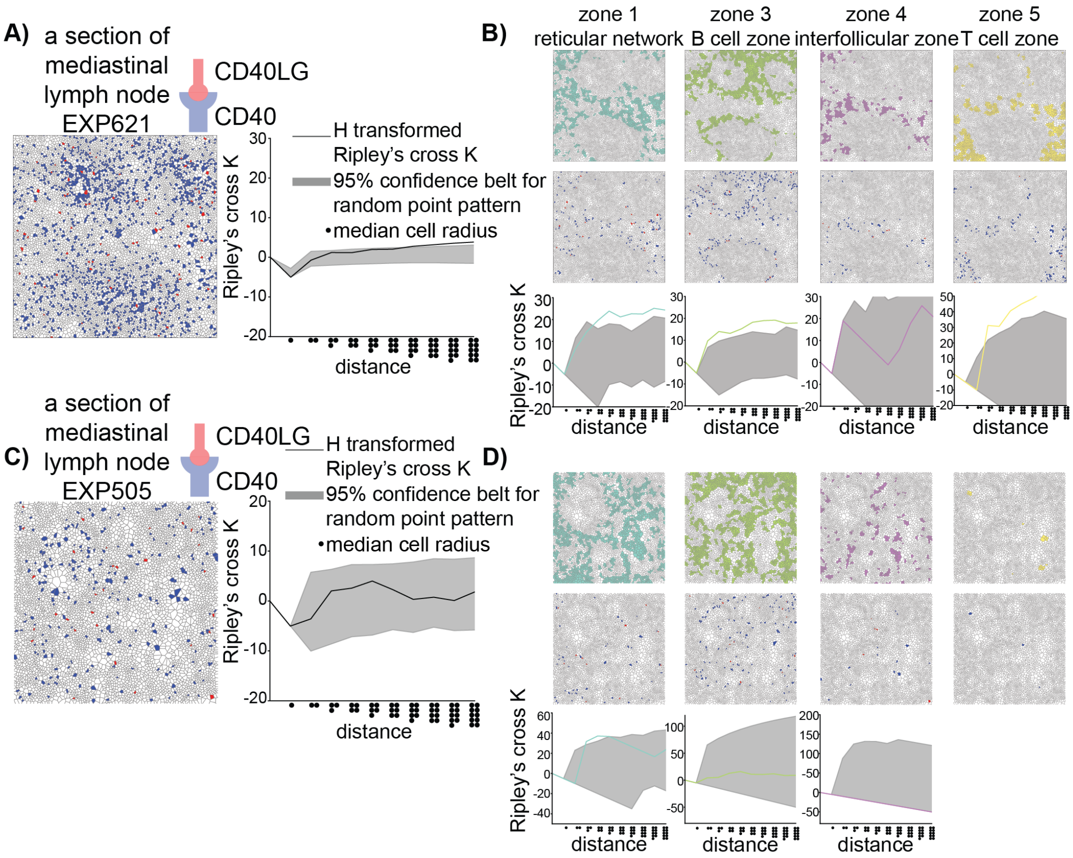


Figure 18: **SPARROW facilitates direct comparison of L-R relationships between two LNs.** **A**: Co-localisation patterns of CD40 positive (receptor, blue) and CD40LG positive (ligand, red) cells in a select 1000 pixel × 1000 pixel window in the mediastinal LN (EXP621) (left). H-transformed Ripley's cross K of CD40 positive cells relative to CD40LG positive cells as a function of increasing distance (black line) (right) in units of median cell radii. 1000 iterations of spatial random patterns were generated by randomly shuffling CD40 positive and CD40LG positive cell labels and Ripley's cross K was calculated for every pattern iteration. The 95% confidence envelope of the random patterns was plotted (gray belt) and compared against the experimental curve. **B**: Zone assignments (top) and within zone co-localisation patterns (middle) of CD40 positive (receptor, blue) and CD40LG positive (ligand, red) cells in the same section as in **A**. H-transformed Ripley's cross K of CD40 positive cells relative to CD40LG positive cells as a function of increasing radii (black line) was calculated for each zone (bottom). 1000 iterations of spatial random patterns were generated by shuffling labels of CD40 positive and CD40LG positive cells within individual microenvironment zones. Zones 1, 3,4 and 5 are shown here and zone 2 is shown in Main Figure 3. **C, D**: Co-localization patterns and Ripley's cross K calculation in the mesenteric LN (EXP505). The methodology is consistent between **B** and **D**. Note that Ripley's cross K calculation was not done for zone 5 in EXP505 as there was no CD40 positive cell in the selected window.
